# Dynamic patterns of correlated activity in the prefrontal cortex encode information about social behavior

**DOI:** 10.1101/2020.08.05.238741

**Authors:** Nicholas A. Frost, Anna Haggart, Vikaas S. Sohal

## Abstract

New technologies have made it possible to measure activity from many neurons simultaneously. Nevertheless, most studies still analyze the activity of simultaneously recorded neurons one-at-a-time, then group together neurons which increase their activity during similar behaviors into an ‘ensemble.’ This notion of an ensemble ignores the ability of neurons to act collectively, and encode and transmit information in ways that are not reflected by their individual activity levels. We used microendoscopic GCaMP imaging to measure prefrontal activity while mice were either alone or engaged in social interaction. We developed new approaches, using neural network classifiers and surrogate (shuffled) datasets, to characterize how neurons synergistically transmit information about social behavior. Surrogate datasets which preserve behaviorally-specific patterns of coactivity (correlations) outperform those which preserve behaviorally-driven changes in activity levels but not correlated activity. This shows that prefrontal neurons act collectively to transmit information about socialization, because social behavior elicits increases in correlated activity that are not explained simply by the activity levels of the underlying neurons. Notably, this ability of correlated activity to enhance the information transmitted by neuronal ensembles is lost in mice lacking the autism-associated gene Shank3. These results show that synergy is an important concept for the coding of social behavior which can be disrupted in disease states, reveal a specific mechanism underlying this synergy (social behavior increases correlated activity within specific ensembles), and outlines methods for studying how neurons within an ensemble can work together to encode information.

## INTRODUCTION

Activity in different neurons often exhibits precise temporal relationships that are modulated by behavior (1–3). For example, subsets of neurons can exhibit correlated activity in which they become co-active at the same time or within short windows of time during specific behaviors. However, correlated activity can occur simply as a byproduct of the interconnected nature of neuronal networks (4). Thus it is unclear whether correlated activity contributes in a meaningful way to information encoding (5), or whether it is simply a reflection of (and thus redundant with) changes in other variables, e.g., firing rates (6).

Many experimental and theoretical studies have studied how noise correlations, e.g., correlations between the trial-to-trial variability of different neurons, affect information coding (7–9). From a population coding perspective, noise correlations can either impair or enhance decoding, depending on whether correlations are present between neurons that have similar or distinct tuning for a behavioral variable of interest. A different question, separate from that of whether variability (noise) between neurons is correlated on a trial-by-trial basis, is whether neurons exhibit correlations that differ based on a behavioral variable. For example, a group of neurons may exhibit correlated activity (co-activity that occurs more often than expected by chance) only during a specific behavior. Groups of co-active neurons represent an attractive computational unit for information processing because they should optimize temporal summation in downstream targets. Thus, even if a group of neurons do not increase their activity levels during a behavioral state, they could increase their correlated activity, and thereby more effectively activate a common downstream target. In this manner, behaviorally-driven changes in correlations can transmit information independent of changes in the activity levels of the underlying neurons. Whereas noise correlations, as their name suggests, reflect correlations in noise, i.e., variability that is unrelated to a behavioral variable of interest, behaviorally-modulated correlations represent potentially important signals in neural networks.

While multiple studies have shown that behavior can modulate correlations (3,10) the functional significance of this has remained unclear, because changes in correlations might simply reflect (and be redundant with) variation in activity levels (6) rather than contributing additional information. Even with the advent of new technologies for simultaneously recording from large numbers of neurons in behaving animals, many studies still focus on information that is transmitted by changes in the activity levels of neurons, while ignoring contributions from correlated activity or other ways in which neurons can act collectively. In particular, it is now commonplace to identify ‘neuronal ensembles’ by grouping together neurons which significantly increase or decrease their activity during the same behaviors or in response to the same stimuli. However, this approach ignores information that is transmitted collectively, e.g., it would not be sensitive to the hypothetical group of neurons described above, which increase their correlations during a specific behavior while keeping their activity levels constant. Thus current methods for identifying neuronal ensembles might falsely conclude that a group of neurons do not encode a behavioral variable (when they do encode it collectively), incorrectly estimate the amount of information that is being encoded, and/or miss important mechanisms that contribute to encoding.

Correlations have been shown to contribute additional information for small groups (3-8 neurons) of cortical neurons (11). Recently, a few studies have examined how correlations contribute to encoding within larger cortical ensembles, which have greater potential for both synergy and redundancy. One found that the identity of a conditioned stimulus was encoded in mean activity levels, but not in moment-to-moment patterns of co-activity (12). In hippocampal region CA1, disrupting correlations impairs the decoding of position, head direction and speed (13). And during a two-choice discrimination task with probabilistic outcomes, correlations between neurons in secondary motor region M2 enhance the encoding of the choice(14). Thus, some studies have found evidence that correlations between neurons can augment the information those neurons encode about behavioral variables(11,13,14). However, all of these focused on the potential role of noise correlations in enhancing encoding by activity levels. I.e., they all hypothesize that noise correlations help disambiguate trial-by-trial variation in activity levels that is related to behavioral variables of interest, from variation that is attributable to ‘noise’, e.g., changes in other variables such as internal states. They did not specifically examine whether correlations might themselves encode unique information by increasing or decreasing in relation to specific behavioral variables, creating behaviorally-specific patterns of co-activity. Thus specific mechanisms through which cortical neurons can collectively encode information currently remain poorly described. A further challenge is the lack of methods for identifying neurons that contribute to the collective encoding of behavioral variables, but do so without significantly increasing or decreasing their activity levels. This makes it difficult to directly examine these neurons in order to assess whether they contribute to ensemble encoding by altering their correlations, generating behaviorally-specific patterns of co-activity, etc.

The goal of this study is to address these two challenges related to neural encoding in the cortex by focusing on a subject that has already been well-studied: the encoding of rodent socialization by neurons in the medial prefrontal cortex (mPFC) (15–18). Many studies have already shown that prefrontal neurons can significantly increase or decrease their activity levels during periods of social interaction (2,17,18) and/or in response to social stimuli such as odors (19). However, these studies have not assessed whether prefrontal neurons might act collectively to encode additional information (beyond what activity levels encode), examined whether this might occur via behaviorally-driven changes in correlated activity, or identified ensembles based on neurons’ ability to encode information collectively, rather than through increases or decreases in their individual activity levels.

The goal of this study is to examine a neuronal population that is already known to encode social behavior, and determine whether this encoding occurs solely via changes in the activity levels of neurons vs. whether additional information is transmitted collectively, and if so, how this occurs. These are critical questions, because if we do not understand the mechanisms which normally encode information, we cannot study how they are disrupted in disease states. Thus by design, we focused on these foundational questions about the nature of information transmission, rather than trying to tackle more nuanced issues, e.g., does this encoding differ between subpopulations of prefrontal neurons, etc.

We used microendoscopic GCAMP imaging in freely-moving mice to measure activity in prefrontal ensembles during periods when mice were either alone or engaged in social behavior. We used a neural network classifier to quantify how well prefrontal ensembles would transmit information about behavior to downstream neurons. By examining the operation of this neural network and using surrogate datasets which preserve activity levels but either preserve or disrupt patterns of correlated activity, we find that behaviorally-driven changes in correlations enhance the information transmitted by neuronal ensembles. Notably, this was not the case in a mouse model of autism (Shank3 knockout mice), even though large numbers of prefrontal neurons are still recruited by socialization in Shank3 knockout mice, and encode socialization through changes in their activity levels. This illustrates that the ability of prefrontal neurons to collectively encode information may be selectively disrupted in pathological states.

## RESULTS

### Social interaction recruits prefrontal ensembles

We implanted microendoscopes (nVoke; Inscopix) into the medial prefrontal cortex (mPFC) of adult wildtype C57/B6 mice (WT) to image calcium transients using GCaMP6f expressed under control of the human synapsin promotor. We imaged freely moving mice during an assay which sequentially introduced 2 novel juvenile mice to the home cage of the subject mouse, first during an initial (novel) epoch and then again during a subsequent (familiar) epoch. These four epochs of social interaction were interleaved with epochs during which the subject mouse was alone in its home cage (‘home cage’ epochs). The first 5 minutes of each interaction epoch was scored by a blinded observer, and each wild-type mouse spent approximately 10 minutes interacting with the juvenile mice (393 +/− 25 s during the novel epochs and 235 +/− 18 s during the familiar epochs, p = 0.00017, paired t test, n = 10 WT mice).

We processed data using a modified PCA/ICA approach (20,21) to identify neurons which were active during the imaging session. To minimize the influence of the surrounding neuropil on neuronal signals, we calculated the mean signal within each ROI, then subtracted the mean signal calculated from a circular annulus surrounding each ROI (Figure S1). Casual inspection of calcium traces revealed that some neurons were more active during epochs of social interaction (compared to periods of home cage exploration), whereas others exhibited the opposite pattern (**Figure 1A**). Correspondingly, aligning calcium traces to the onset of social interaction revealed many neurons that either increased or decreased their activity levels at the onset of interaction (**Figure 1B**). Fluorescence traces were converted to binary event rasters (see Methods for details of event detection), in which most neurons were “active” in less than 5% of frames (**Figure 1C**). As a population, imaged neurons were more active during social interaction (**Figure 1C**, n = 663 neurons from 10 mice, percent time active in home cage: 1.8% +/− 0.1, percent time active during social interaction: 2.1 +/− 0.1%, p = 0.00002, paired t-test). There was a bimodal distribution of cells that were significantly more (>90^th^ percentile, social: 152/663 neurons, home cage: 80/663 neurons; p < 0.00001, Chi-Squared Test) or less active (<10^th^ percentile, social: 128/663 neurons, home cage: 119/663 neurons; p = 0.5) during either social interaction or matched periods when mice were alone in their home cage, as compared to circularly shuffled datasets (**Figure 1D**). These correspond to conventionally-defined neuronal ensembles, i.e., groups of neurons that increased or decreased their activity levels during social interaction.

**Figure 1.**
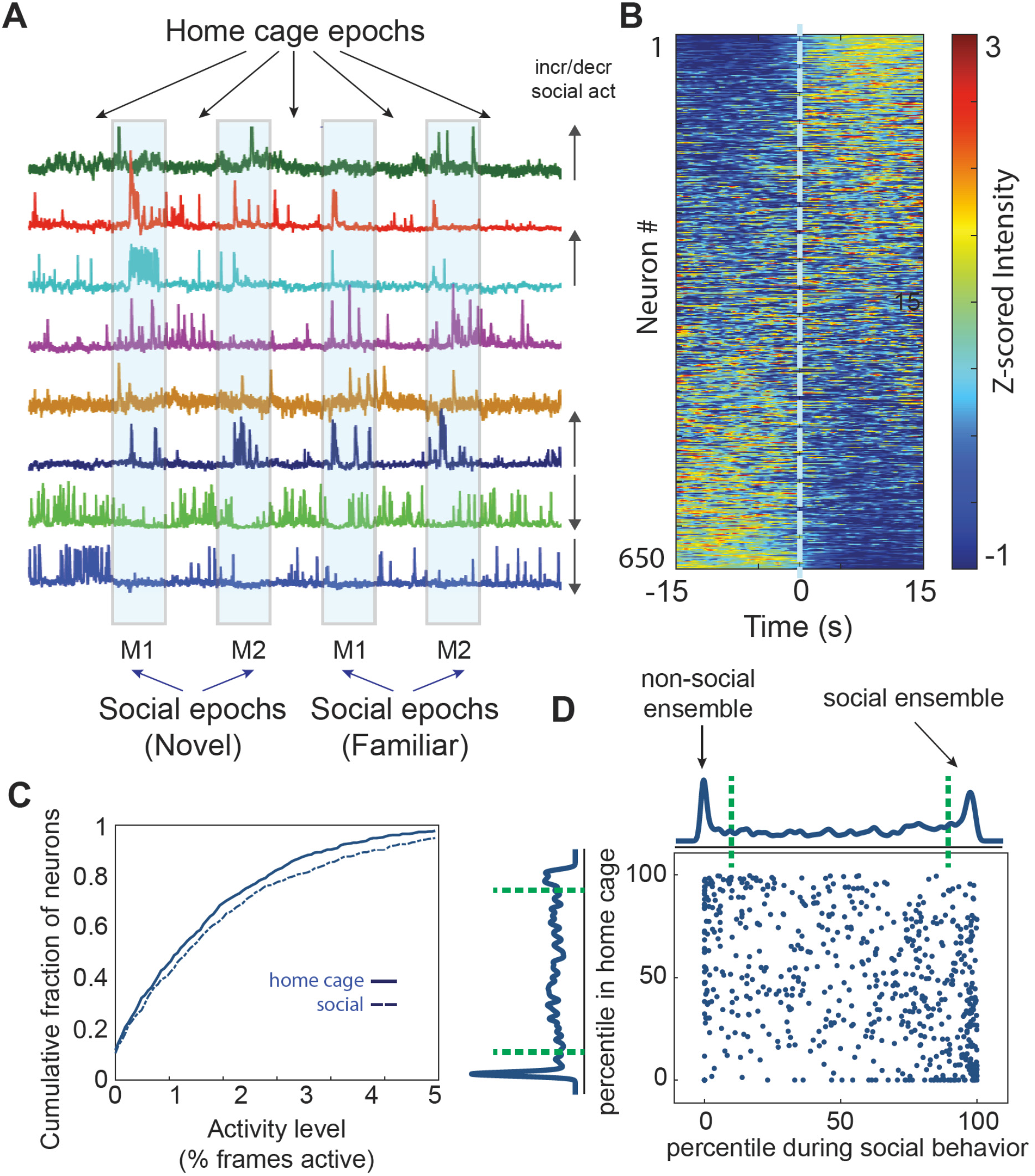
Social interaction modulates activity levels within prefrontal ensembles. **A.** Mice were imaged across 9 consecutive behavioral epochs (each lasting 5 min) during which they were either alone in their homecage or interacted with one of two novel sex-matched juvenile mice introduced to the homecage (‘M1’ or ‘(M2’). Each novel mouse was subsequently re-introduced to the home cage during a familiar epoch. GCaMP traces during show examples of neurons that appear to increase or decrease activity during social epochs (see arrows at the right of each trace). **B.** Mean z-scored GCaMP traces for all neurons recorded from wild-type mice (663 neurons from 10 mice) aligned to the onset of social interaction during the first bout of interaction within each social epoch. **C.** Cumulative plot showing the distribution of activity levels for individual neurons during homecage epochs or periods of social interaction (percent time active in homecage: 1.8% +/− 0.1, percent time active during social interaction: 2.1 +/− 0.1%, p = 0.00002, paired t-test; n = 663 neurons from 10 WT mice). **D.** Scatter-plot showing the activity of each neuron during each behavioral condition, expressed as a percentile relative to a null distribution generated by circularly shuffling that neuron’s activity. Activity levels during social interaction or while the mouse was alone in its home cage are plotted on the x and y axis, respectively. Kernel density plots along the axes indicate the fraction of neurons whose activity was at a given percentile of the null distribution. Neurons with activity > 90^th^ percentile of shuffled datasets (green dotted line) were considered to be positively modulated, whereas neurons with activity < 10^th^ percentile (green dotted line) were considered to be negatively modulated during each behavior (>90^th^ percentile, social: 152/663 neurons, home cage: 80/663 neurons; p < 0.00001, chi-squared test; <10^th^ percentile, social: 128/663 neurons, home cage: 119/663 neurons; p = 0.5, chi-squared test).

### Using a neural network classifier to assess how well ensembles transmit information

Next, we sought to determine how well these prefrontal ensembles would transmit information about social behavior to downstream neurons. Specifically, we measured how well downstream neurons could decode whether a mouse was engaged in social behavior based on input from these prefrontal neurons. Later we will study how this was altered in Shank3 knockout mice. For this we used a simple neural network classifier that received input from the recorded neurons. Our rationale for using this kind of neural network classifier was threefold. First, a simple neural network measures information that is immediately and readily available to downstream neurons. Second, for a neural network with only one hidden layer, it is straightforward to examine the weights to determine how the network performs the classification. This can provide insight into exactly how the neural network is able to decode behavior from the input activity. Third, examining how various parameters of the neural network affect its performance can provide additional clues about how information is represented within the input population.

**Figure 2A** shows the design of the neural network classifier. For clarity, we use the terms ‘neurons’ specifically to refer to actual prefrontal neurons (which provide input to the neural network), and ‘units’ to refer to simulated elements within the network. The network consisted of a hidden layer containing 1000 units. We chose this number because it is both an order of magnitude larger than the number of input neurons and an order of magnitude smaller than the number of frames available for training (the latter helps ensure that there will be enough data to train the output weights). We simulated a different neural network for each mouse. Each hidden layer unit received input from a random subset of the prefrontal neurons from one mouse. I.e., each frame represents one timepoint and if neuron *i* is active in a frame then it provided an input of 1 to all the hidden units to which it is connected; otherwise it provides an input of 0. For each simulation, there was a fixed connection probability between each input neuron and each hidden layer unit. We tried different values for this connection probability in order to measure how classifier performance depends on the number of neurons that provide input to each hidden layer unit. Each hidden layer unit had an output weight which specifies how strongly that unit excites or inhibits a single output unit which classifies activity as belonging to periods in which a mouse was actively engaged in social interaction or alone in its home cage. E.g., output unit activity < 0.5 corresponds to the social condition, while output unit activity > 0.5 corresponds to the home cage condition. These output weights were adjusted during training (see Methods for details of the training rule) while the pattern of input connectivity was fixed. This models the situation in which prefrontal neurons transmit information to a downstream population of neurons (the hidden layer) that decode behavior via their output weights. We initially trained networks on 50% of the data (frames) and used the held-out data for testing. We trained and tested using intervals during which the mouse was actively engaged in social interaction (i.e. actively sniffing / nose in contact with subject mouse) or matched intervals when the mouse was alone in its home cage.

**Figure 2.**
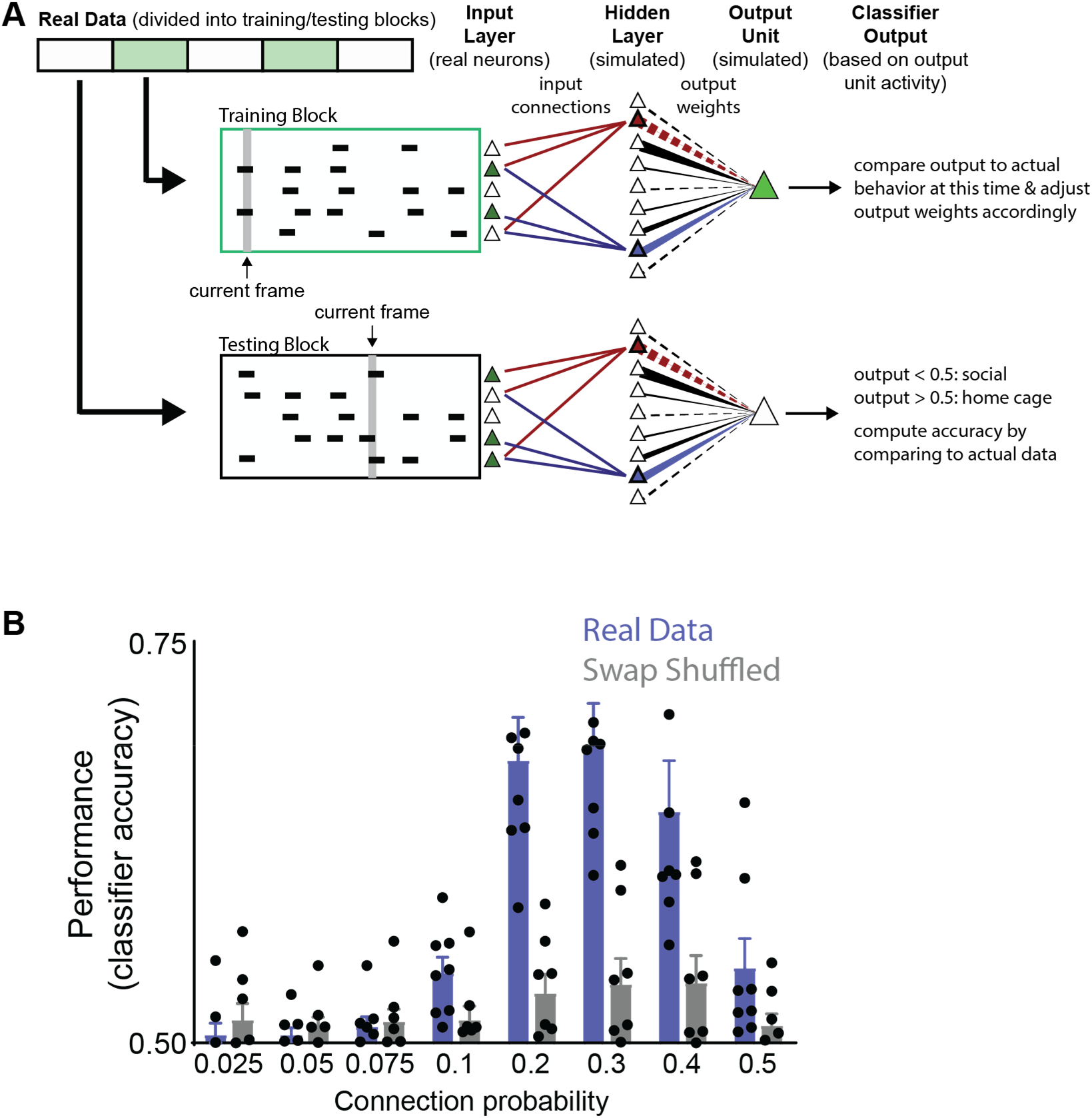
Classifying behavior from prefrontal ensembles using a simple neural network. **A.** We constructed a neural network consisting of a single hidden layer (containing 1000 units) which were connected to a single output unit. The thickness of lines between each hidden layer unit and the output unit reflects the magnitude of the output weight. Positive and negative weights are indicated by solid and dashed lines, respectively. Each hidden layer unit received input from a random subset of prefrontal neurons from one real dataset. For clarity, we have only shown input connections to two hidden layer units (which are differentiated by their blue and red colors) – output weights from other hidden units are shown in black. The output weight from each hidden layer neuron was iteratively updated during training. We trained the classifier to distinguish periods marked as home cage exploration or social interaction by dividing a dataset into 500-frame blocks, and then using alternating blocks for training or testing. **B.** The classifier performed poorly (near chance) when the input connection probability (governing the number of prefrontal neurons that provided input to each hidden layer unit) was <10%. Classification accuracy was above chance in 8/10 mice and increased to a peak of 69 +/− 3% in these mice, before decreasing again for connection probabilities >30%. The classifier performed near chance levels when we trained and tested using data that had been randomly swap-shuffled.

Because each hidden unit simply sums the activity of a fixed population of mPFC neurons, one can think of each hidden unit as representing one ensemble of mPFC neurons. The size of these ensembles is determined by the input connection probability. In this framework, training serves to determine how well each ensemble (corresponding to an individual hidden unit) is correlated with behavior. Hidden units corresponding to ensembles which are strongly associated with one type of behavior will be assigned large positive or negative output weights, whereas those that are not well correlated with behavior will have output weights near zero.

Critically, we did not attempt to classify more nuanced behaviors, besides whether the subject mouse was exploring another mouse vs. alone in its home cage. This was by design because doing so would have reduced the number of training frames available for each behavioral label. A large number of training frames for each behavioral label are necessary to train a neural network with a large number of hidden units, and a large number of hidden units are necessary in order to effectively capture possible collective encoding (because, as noted above, each hidden unit effectively represents one neuronal ensemble). Thus, subdividing periods of social interaction into more nuanced behavior would have compromised our ability to resolve collective encoding. Therefore, because the focus of this paper is on trying to study collective encoding, we intentionally used simple, binary classifications of behavior (e.g., social vs. home cage).

### Classifier performance is optimal for intermediate connection probabilities

Classifier performance was strongly dependent on the probability that each input neuron was connected to each hidden unit. For the 8/10 datasets that could be classified above chance, classifier performance (measured on the 50% of data which was held-out during training) was near chance levels when the connection probability was < 0.1, but increased to a peak of 69 +/− 3% (**Figure 2B**; **Table S1**) for a connection probability of 0.3. Accuracy decreased dramatically when the connection probability increased to 0.5 indicating that connection probabilities ~0.2 - 0.4 are optimal.

We also validated classifier performance by training and testing on surrogate datasets that were generated by ‘swap shuffling’ our original datasets. We created ‘swap shuffled’ surrogate datasets by randomly swapping blocks of activity between neurons (each block of activity = a set of consecutive frames during which the neuron was active). To understand this, think of the entire raster as a collection of blocks of activity. Each block occurs at a specific time, has a specific duration, and is associated with a particular neuron. Swap shuffling is equivalent to just shuffling the neurons associated with each block of activity (the start time and duration of each block do not change). For example, if neuron *i* originally became active at time *t1* for *n1* frames and neuron *j* was active at time *t2* for *n2* frames, then in the surrogate dataset neuron *i* might become active at *t2* (but not at *t1*) while neuron *j* might become active at *t1* (but not *t2*). Swap shuffling preserves the number of neurons active at each point in time (because the timing of blocks of activity does not change). It also preserves the number of blocks of activity for each neuron, and this tends to preserve the overall level of activity of each neuron. Activity levels are not perfectly preserved, because blocks of activity can have different durations. Nevertheless, in practice, blocks of activity tend to have similar durations and the similarity between the mean activity level in each neuron before and after swap shuffling of entire datasets was 0.97 +/− 0.01. As expected, we found that neural network classifiers trained and tested on swap shuffled datasets performed near chance levels (Figure 2B).

### Prefrontal neurons that drive classifier performance change their correlations during social behavior

Next, we examined connections in trained networks to reveal factors which enable them to successfully classify social vs. home cage behavior (we analyzed networks with a connection probability = 0.3 since this maximized performance of the population). Each hidden layer unit has an output weight which measures how strongly it excites or inhibits the output unit that represents the “decision” (social vs. home cage). Hidden units with output weights ~0 don’t contribute to this decision. By contrast, hidden units with strong negative or positive weights promote the social or home cage decision, respectively (Figure 3A). Therefore, we hypothesized that there might be important differences in the pattern of input to hidden units, depending on whether those hidden units have large positive or negative output weights.

**Figure 3.**
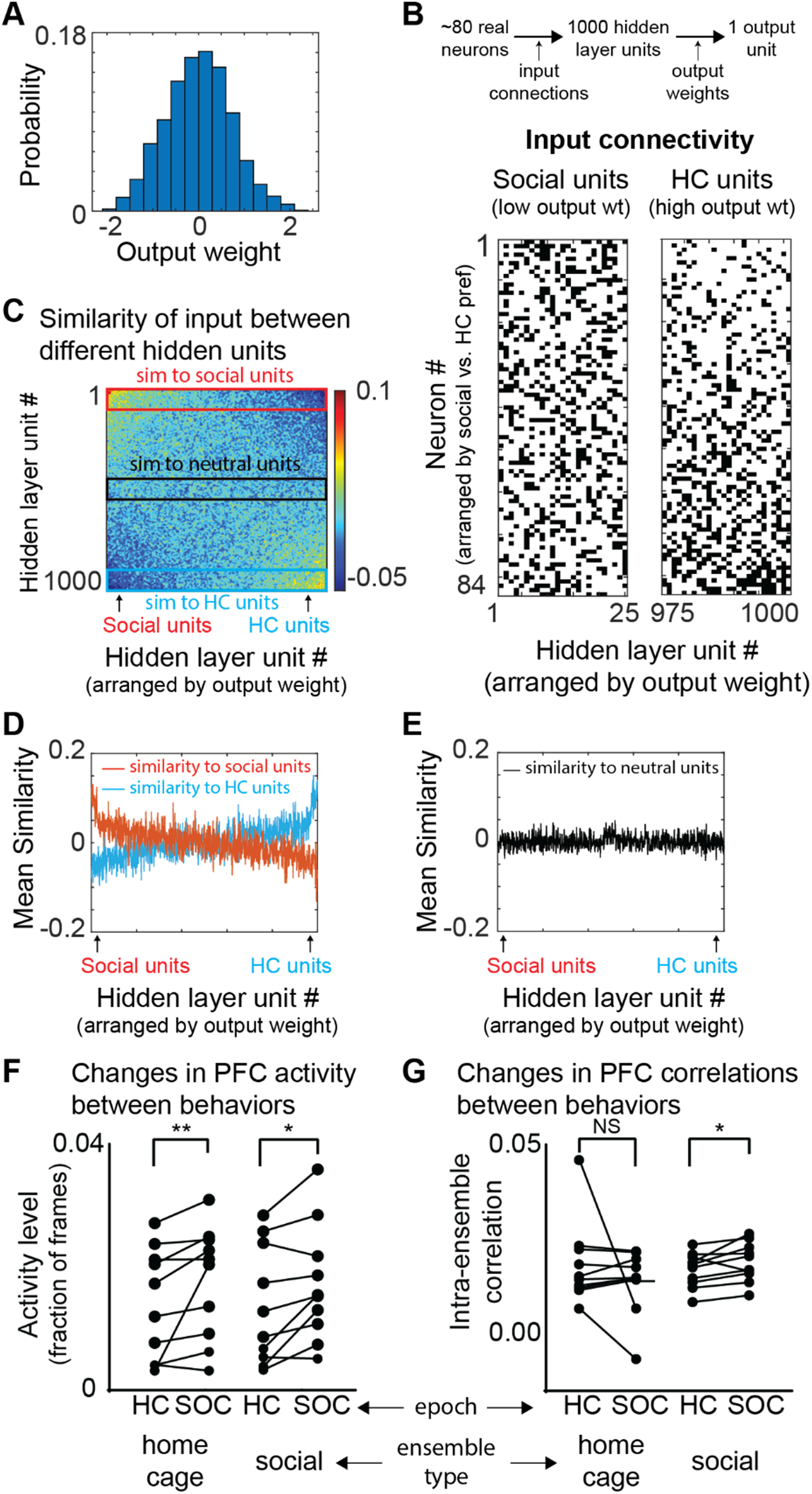
Classifier weights reveals an ensemble that increases correlations during social behavior and detects social behavior. **A.** Example histogram depicting the distribution of output weights assigned to connections between hidden layer units and the output unit over the course of training. **B.** Matrix of input connections for hidden units which detect the social (left) or home cage condition (right). The hidden layer units (x-axis) have been arranged in order of increasing output weights to identify ‘social units’ (25 most negative output weights) and ‘home cage units’ (25 largest positive output weights). Prefrontal neurons (y-axis) have been arranged in order of their preference for social interaction vs. home cage, i.e. the difference between their activity levels in the two conditions. **C.** Correlation matrix showing the input similarity, i.e., the pairwise correlation between binary vectors representing the input connections to each pair of hidden layer units. Hidden layer units are arranged in order of increasing output weight. Red and blue rectangles indicate correlations with social or home cage units, respectively. A gaussian filter with a standard deviation of 3 was applied to the 1000×1000 matrix to improve visualization. **D.** For each hidden layer unit, we plotted its average input similarity to either the 25 social units (red) or the 25 home cage units (blue). Hidden layer units (x-axis) are again arranged by output weight. Social units had similar patterns of input compared to each other but not to home cage units and vice-versa. **E.** The average input similarity of each hidden layer unit to 25 hidden layer units with near-zero output weights (‘neutral units’; black rectangle in C). **F.** We defined social and home-cage (HC) ensembles as the 20% of prefrontal neurons most likely to provide input to the social or home cage units, respectively. The mean activity of both home cage and social ensembles increased during social interaction compared to the home cage condition (social ensemble: mean activity level 1.4 +/− 0.3% in home cage vs. 1.8 +/− 0.3% during social interaction, p < 0.05, sign-rank test; home cage ensemble: mean activity level 1.5 +/− 0.30% in home cage vs. 1.9 +/− 0.3% during interaction, p < 0.001, sign-rank test). **G.** Correlations between neurons in the same ensemble increased during social interaction for the social ensemble but for the home cage ensemble (mean correlation coefficient between neurons in the social ensemble: 0.009 +/− 0.002 in home cage vs. 0.012 +/− 0.002 during social interaction, p < 0.05; home cage ensemble mean correlation coefficient 0.011 +/− 0.02 in home cage vs. 0.005+/− 0.003 during social interaction, p=0.99, sign-rank).

We arranged hidden layer units based on their output weights, i.e., the unit with the most negative weight was unit 1 and the unit with the most positive weight was unit 1000. Then we defined the 25 hidden layer units with the most negative weights as “social units” and the 25 with the most positive weights as ‘home cage units’ (**Figure 3B**). For comparison we also defined the 25 hidden layer units with the weights closest to zero as ‘neutral units.’ For each pair of hidden units, we computed the similarity between their inputs (i.e., the correlation between their input vectors; **Figure 3C**). We then plotted the average input similarity of each hidden unit to either the social or home cage units (**Figure 3D**) or the neutral units (**Figure 3E**). Social and home cage units tended to receive input from the same prefrontal neurons as other hidden layer units with the same preference, i.e., which also had negative or positive output weights. By contrast neutral units did not exhibit any such relationship.

The preceding suggests that distinct ensembles of prefrontal neurons provide input to either social or home cage hidden units. We hypothesized that there might be important features of activity in these ensembles that support the classification of social vs. home cage behavior. For example, one possibility is that prefrontal neurons which provide input to social units might tend to increase activity during social behavior, whereas prefrontal neurons which provide input to home cage units do the opposite. Surprisingly, this was not the case. In fact, both ensembles of prefrontal neurons significantly increased their activity when mice were engaged in social interaction (**Figure 3F**; social ensemble: mean activity level 1.4 +/− 0.3% in home cage vs. 1.8 +/− 0.3% during social interaction, p < 0.05, sign-rank test; home cage ensemble: mean activity level 1.5 +/− 0.3% in home cage vs. 1.9 +/− 0.3% during social interaction, p < 0.001, sign-rank test). Reasoning that changes in correlations would alter the frequency with which pairs or groups of neurons became co-active, we examined pairwise correlations between the activity of prefrontal neurons within each ensemble. Strikingly, mean correlations within the social ensemble increased during social interaction (**Figure 3G**; (mean correlation coefficient between neurons in the social ensemble: 0.009 +/− 0.002 in home cage vs. 0.012 +/− 0.002 during social interaction, p < 0.05). By contrast, there was a non-significant decrease in correlations within the home cage ensemble (Figure 3G; home cage ensemble mean correlation coefficient 0.011 +/− 0.02 in home cage vs. 0.005 +/− 0.003 during social interaction, p=0.99, sign-rank).

Thus, the ensemble of prefrontal neurons which provide input to the social units form an assembly that collectively becomes more co-active (correlated) during social behavior. In contrast, the prefrontal neurons which provide input to the home cage units increase their activity, but not their co-activity, during social behavior. This suggests that behaviorally-driven changes in correlations may contribute to the encoding of social behavior.

### Swap shuffling within a behavioral state preserves information encoded by activity levels but disrupts encoding by correlated activity

How can we quantitatively assess the contribution of these behaviorally-modulated correlations to classifier performance? Ideally we would first train a neural network on the original data. Then we would test this network’s ability to classify data which maintained behaviorally-driven changes in activity levels, but either removed or preserved the specific patterns of correlated activity found in the original dataset. Indeed, we have already developed methods for shuffling that achieve these goals. First, to shuffle the data in a manner that maintains behaviorally-driven changes in activity levels, but disrupts the patterns of correlated activity, we can swap shuffle activity, but do so within each behavioral condition rather than across the entire testing dataset. In other words, we first divide up the raster into separate subrasters for each 5 minute behavior epoch (when the mouse was either engaged in social interaction or alone in its home cage). Then we performed swap shuffling (as described above) separately on each subraster, before recombining these swap shuffled subrasters to create the swap shuffled surrogate dataset for testing. Because swap shuffling tends to preserve activity levels, and because we swap shuffled activity within a behavioral condition, neurons that increase or decrease activity during periods of social interaction in the original dataset will also do so in the swap shuffled surrogate dataset.

We first confirmed that swap shuffling preserves information encoded by activity levels, while disrupting the ability of correlated activity to transmit information. For this we created synthetic datasets, swap shuffled them, and then measured classifier performance (**Figure 4**). Specifically, we generated different types of synthetic datasets, which encoded two different ‘behavioral states’ (state A or state B) via correlations or activity levels. In all cases, we first created a raster for ‘state A’ by randomly assigning the activity of 100 neurons so that the total activity in each frame oscillated around a mean level. To encode ‘state B’ using correlations, we started with the original ‘state A’ raster, but shifted the activity of individual neurons in time, to create between 1 and 5 cell assemblies composed of co-active neurons. Specifically, each assembly contained 8 neurons; whenever one of these neurons was active, we probabilistically shifted the activity of other neurons in that assembly, so that they would be co-active in the same frame (**Figure 4A**). Notably, the activity level of every neuron was the same in the original ‘state A’ raster, and these newly created ‘state B’ rasters. State B was distinguished simply by an increase in correlated activity. (When we shifted the activity of one neuron in an assembly into a frame, we swapped its activity with that of another neuron that was outside of the assembly; as a result, the total activity in each frame was unchanged).

**Figure 4.**
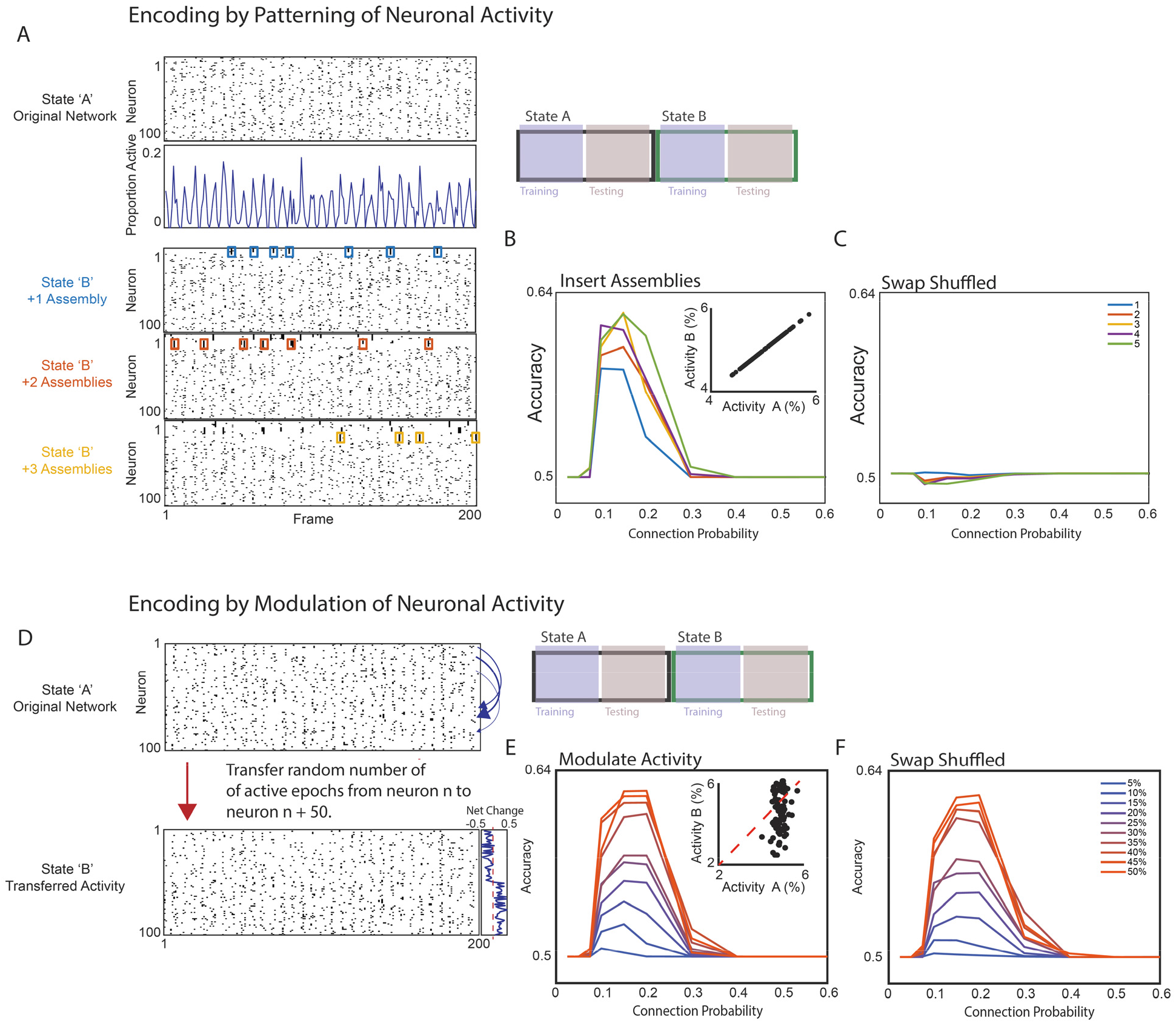
Quantification of state-dependent patterned activity with a neural network-based classifier. **A.** We generated synthetic datasets consisting of 100 neurons. Each dataset consisted of two activity rasters, corresponding to states ‘A’ and ‘B’. Each raster contained 6,000 timepoints (frames). The overall level of activity in each raster oscillated around a mean level of 5% (e.g., 5% of neurons active in a frame). In ‘State A’ neuronal activity was randomly assigned. Then we created 1-5 cell assemblies in ‘State B’ by rearranging the activity raster for ‘State A’. Specifically whenever the first neuron in an assembly was active, we swapped activity in the rest of the network, so that other neurons in the same assembly would be co-active in the same frame. This was achieved by making reciprocal swaps between neurons. E.g., suppose a neuron outside the assembly was active in the desired frame. Then we swapped its activity with a neuron within the assembly that was active in a different frame. Thus neither the total number of neurons active in a given frame, nor the total activity level of any neuron differed between the ‘State A’ and ‘State B’ rasters. As a result, ‘State B’ was differentiated from ‘State A’ only by 1, 2, 3, 4, or 5 assemblies comprised of correlated (co-active) neurons. **B.** A sparsely connected binary classifier perfomed above chance in distinguishing ‘State A’ from ‘State B’ when 50% of the data was used for training, and the remainder for testing. Inset shows the activity level of each neuron in ‘State A’ plotted against its activity level in ‘State B.’ **C.** The classifier, trained on the original ‘State A’ and ‘State B’ rasters, performed at chance levels when tested on swap-shuffled versions of the ‘State A’ and ‘State B’ rasters. **D.** We again generated synthetic datasets. The ‘State A’ rasters were as described in panel *A*. ‘State B’ rasters were generated by transferring a proportion of the activity from the first 50 neurons in each State A raster to neurons 51-100. In this manner, half the neurons increased their activity in ‘State B’ compared to ‘State A’, while the other half decreased their activity in ‘State B’ compared to ‘State A.’ The proportion of activity transferred was determined by randomly drawing a number with an upper bound ranging between 5 and 50%. **E.** We used 50% of the data to train a sparsely connected binary classifier to distinguish ‘State A’ from ‘State B’. The remaining data was used for testing. Insert shows the activity level of each neuron in ‘State A’ plotted against its activity in ‘State B.’ **F.** Contrary to panel *C*, swap-shuffling these datasets (which differ in the activity levels of individual neurons) does not disrupt classification accuracy.

After training, our neural network classifiers were able to distinguish between ‘state A’ activity and ‘state B’ activity (**Figure 4B**). This confirms that this type of neural network can classify two different behaviors even when they are encoded entirely by changes in correlated activity, not by any differences in activity levels. Furthermore, when we swap shuffled the ‘state B’ raster to eliminate these correlations, classifier performance was reduced to chance levels (**Figure 4C**).

Next tested the ability of our neural networks to classify ‘state A’ vs. ‘state B’ utilizing differences in activity levels. For this, we again started with a random ‘state A’ raster. Then, to generate a ‘state B’ raster, we shifted a proportion of activity from neurons 1-50 to neurons 51-100 (**Figure 4D**). Thus, in these synthetic datasets, neurons 1-50 have higher activity in ‘state A’ whereas neurons 51-100 have higher activity during ‘state B’. We varied the proportion of activity that was transferred between neurons. Once again, after training, our neural networks could classify activity patterns corresponding to ‘state A’ vs. ‘state B’ (**Figure 4E**). However, in this case, swap shuffling ‘state B’ rasters did not disrupt classifier performance (**Figure 4F**). This confirms that our neural networks classifiers can detect encoding that is based on either activity levels or correlated activity. Swap shuffling a dataset (within one behavioral state) will remove information encoded by correlated activity without diminishing encoding via activity levels.

### SHARC shuffling preserves patterns of correlated activity

Before applying this same approach to our actual datasets, we also wanted to create a method for shuffling that preserves patterns of correlated activity. For this, we used a method that we published previously: SHuffling Activity to Preserve Correlations, or SHARC (22). Like swap shuffling, SHARC also re-assigns blocks of activity between neurons, but rather than doing so randomly, it instead follows an algorithm that achieves a target correlation matrix (in this case, the original correlation matrix) (Figure 5B-C). The full details of SHARC are presented in the Methods. Briefly: on each iteration, we randomly select one block of activity to be assigned to a new neuron. Instead of choosing the new neuron randomly, we first compute the difference between the target correlation matrix and the correlation matrix of the partially reconstructed surrogate dataset. Then we assign the block of activity to the neuron which will optimally reduce this difference. Finally, to maintain the mean activity level of each neuron, there is also an absolute limit on how many blocks of activity can be re-assigned to each neuron. Notably, when we applied SHARC shuffling to our previously described synthetic datasets, it visibly preserved assemblies of coactive neurons, and did not disrupt the ability of neural network classifiers to distinguish between “state A” and “state B” based on either activity levels or correlated activity (Figure S2).

**Figure 5.**
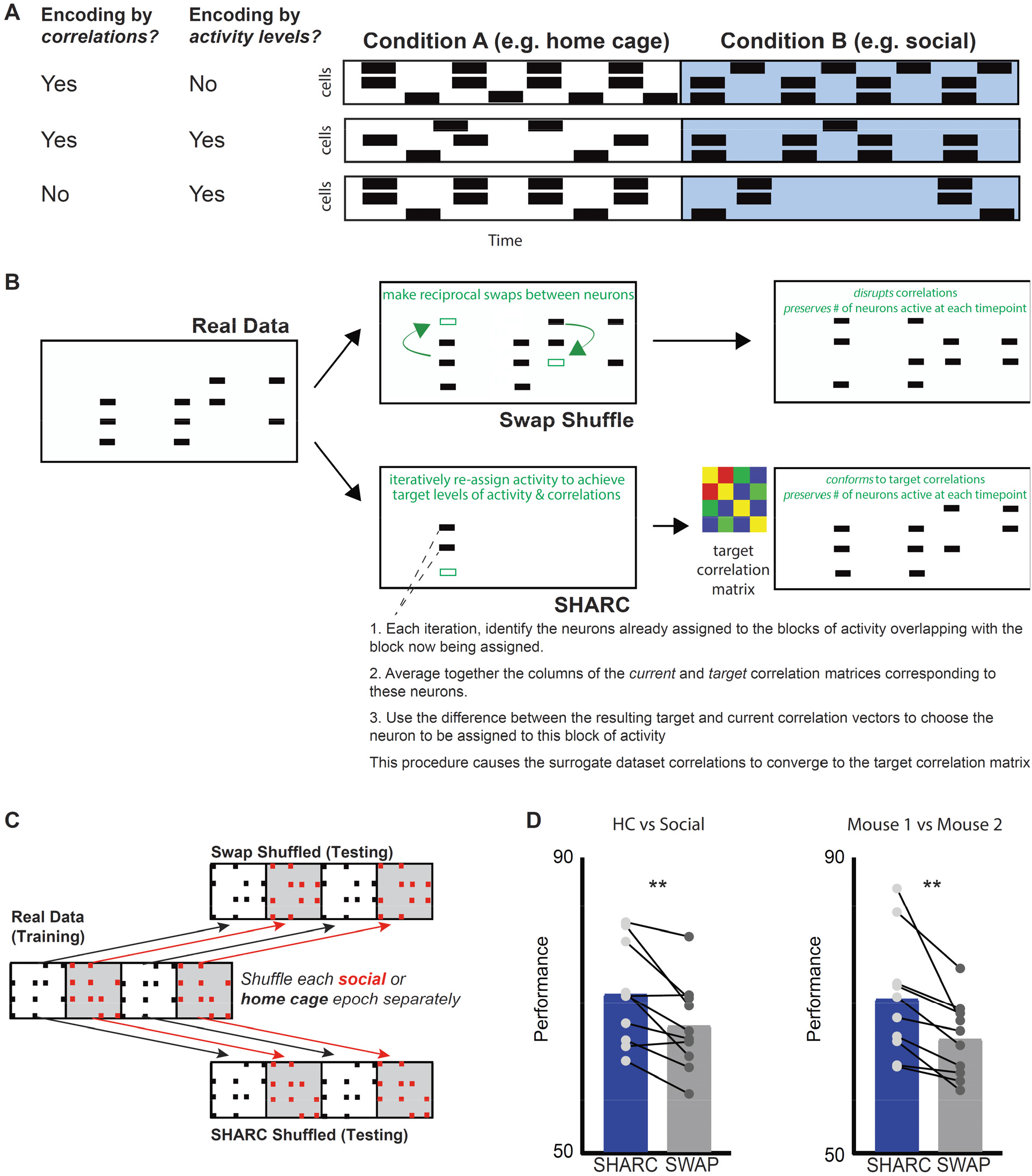
Correlations transmit additional information that is not efficiently conveyed by changes in activity levels alone. **A.** Cartoon illustrating that information about behavior may be encoded through changes in activity levels, correlations between neurons, or both. When behavior modulates activity levels, correlations in two behavioral conditions may differ or be the same, and vice-versa. **B.** To disentangle the roles of activity levels and correlations in transmitting information we used two different methods to create shuffled (surrogate) datasets which preserve changes in activity levels, but either do or do not preserve patterns of correlations. We made random, reciprocal swaps of activity between neurons to generate surrogate datasets which maintained network activity in each frame as well as the number of blocks of activity for each neuron. However, these datasets destroyed the correlation structure. In a second set of surrogate datasets we used SHARC to iteratively generate surrogates in which the correlation structure was also maintained. **C.** To maintain dynamic changes in activity levels and correlations that are associated with the two behavioral conditions we swap-shuffled or performed SHARC separately for each behavioral epoch, then concatenated the 9 resulting surrogate subrasters to create each surrogate dataset. **D.** We trained a classifier (with a connection probability = 0.3) on each real dataset, then tested that classifier on swap or SHARC-shuffled surrogate datasets generated from that real dataset. Accuracy was significantly higher for the SHARC-shuffled surrogates, which maintain the behaviorally-modulated correlations found in the original dataset. Left: accuracy for surrogate datasets in classifying periods of home cage (HC) vs. social (Soc) behavior = 71 +/− 2 (SHARC) vs. 67 +/− 2% (swap); p = 0.005, paired t test. Right: accuracy for surrogate datasets in classifying interaction with Mouse 1 vs. interaction with Mouse 2: 71 +/− 3% (SHARC) vs. 65 +/− 2 % (swap); p = 0.007, paired t test. n = 10 mice.

### Patterns of correlated activity contribute to classifier performance

Having confirmed that we can swap or SHARC shuffle individual activity during individual epochs to preserve activity levels while either disrupting or preserving correlated activity, we now used this approach to quantitatively measure the encoding of social behavior by activity levels vs. correlated activity in our actual datasets. Specifically, we swap or SHARC shuffled each social or home cage subraster separately. Then we concatenated these shuffled subrasters to create swap or SHARC shuffled surrogate datasets that preserve behaviorally-specific levels of activity, while disrupting or preserving patterns of correlated activity, respectively.

Both swap and SHARC shuffled surrogate datasets preserved levels of activity observed during either social interaction or periods when mice were alone in their home cages. Specifically, we computed the correlation between vectors in which each element represents the activity level of one neuron during one behavioral condition, and quantified the correlation between each real and surrogate dataset. For swap shuffled surrogate datasets, the similarity of activity levels (compared to real data) was 0.89 +/− 0.02 in the home cage and 0.82 +/− 0.04 during social interaction. For SHARC shuffled surrogate datasets, the similarity of activity levels (compared to real data) was 0.88 +/− 0.03 in the home cage and 0.86 +/− 0.03 during social interaction (n = 10 mice/datasets). We also computed the similarity of the pattern of correlations between each surrogate dataset and the corresponding real dataset. In this case, only SHARC shuffled surrogate datasets preserved patterns of correlations. For swap shuffled surrogate datasets, the similarity of correlations to the real data was 0.01 +/− 0.01 in the home cage, and 0.03 +/− 0.01 during social interaction. For SHARC shuffled surrogate datasets, the similarity was 0.50 +/− 0.05 in home cage and 0.55 +/− 0.03 during social interaction.

We then trained classifiers on each dataset and tested each classifier using either swap or SHARC shuffled surrogate datasets generated from the same dataset using for training (Figure 5C). Classifiers performed better than chance when tested with either surrogate dataset indicating that changes in activity levels encode behavioral information, making it possible to distinguish bouts of social interaction from periods when a mouse is alone in its home cage. However, classifier accuracy was significantly higher for SHARC shuffled surrogates datasets than for swap shuffled ones (Figure 5D, left; accuracy for classifying home cage vs. social = 71 +/− 2% for SHARC vs. 67 +/− 2% for swap shuffled datasets, p = 0.005, paired t test).

### Patterns of correlated activity also help encode mouse identity

We wondered whether we could use this approach to study how correlations contribute to encoding of additional information besides just whether or not a mouse was engaged in social interaction. In our behavioral design, the subject mouse interacted during one mouse for two epochs and with a second mouse during two different epochs. Therefore, we also trained neural networks to classify whether the subject mouse was interacting with Mouse 1 vs. Mouse 2. Again, a neural network classifier was able to perform this classification above chance levels, and performance was higher when the classifier was tested using SHARC shuffled surrogate datasets than when tested using swap shuffled surrogate datasets (Figure 5D, right: accuracy for classifying Mouse 1 vs. Mouse 2 = 71 +/− 3% for SHARC vs 65 +/− 2% for swap shuffled datasets, p = 0.007, paired t test). This shows that behaviorally-modulated patterns of correlated activity transmit additional information, beyond what is readily decodable from activity levels alone.

### Combinations of coactive neurons occur in a behaviorally-specific manner

Interestingly, neural networks perform classification better for connection probabilities ~0.2 – 0.4 than for connection probabilities < 0.1. When the connection probability is low, each hidden unit receives input from individual prefrontal neurons or small groups of neurons. By contrast, when the input probability is higher, hidden units receive input from larger groups of prefrontal neurons. This suggests that training proceeds more efficiently when the network represents information about social vs. home cage behavior using multineuron combinations, instead of activity within individual neurons or small groups. Together with the fact that classifier performance was higher for SHARC shuffled datasets than swap shuffled ones, this indicates that multineuron patterns of coactivity, rather than just levels of activity within neuronal ensembles, transmit information about social behavior. Therefore as a proof-of-concept, we directly examined whether 3-neuron patterns of coactivity occur in a behaviorally-specific manner. We examined 3-neuron combinations because they measure network structure beyond pairwise correlations and are the building blocks of larger combinations. One could in principle analyze larger combinations, but because of the limited numbers of neurons and frames in our datasets, there is not always adequate statistical power to resolve larger combinations, i.e., to detect large numbers of combinations that occur more often in real datasets than expected by chance.

First, we quantified how often each possible 3-neuron combination occurred in real datasets. Then we calculated how often each of these combinations occurred in datasets that had been swap-shuffled (across the entirety of the dataset). For each real dataset we constructed 1000 swap-shuffled datasets, and identified ‘enriched combinations,’ which occurred more often in real datasets than in 95% of swap shuffled surrogate datasets. Enriched combinations are those which occur more often in real datasets than expected based on the chance overlap of activity between marginally independent neurons. Finally, we quantified how many of these enriched combinations were behaviorally-specific, i.e., occurred exclusively during social or home cage epochs. Combinations could appear to be behaviorally-specific simply because they only occurred at a single timepoint. Therefore we also restricted our analysis to enriched combinations which occurred during multiple distinct bouts of social interaction and/or matched sets of intervals during home cage epochs. Many of these repetitively-occurring enriched combinations were behaviorally-specific: 43.5% occurred during social interaction, 26.5% during home cage epochs, and 30% during both conditions.

The selective occurrence of enriched combinations either during social interaction or when a mouse is alone in its home cage may reflect changes in single neuron activity (i.e., neurons that form a social combination are only active during the social condition), and/or changes in correlations (i.e., neurons are active in both conditions but only *co-active* during social behavior). To test the hypothesis that changes in correlations underlie the behavioral specificity of significant combinations, we examined the 3-neuron combinations that were specifically enriched during either periods of home cage exploration or social interaction (**Figure 6**). We defined specific enrichment as those combinations which occurred more often in real data than in 95% of swap-shuffled surrogate datasets for one behavioral context, and less in real data than in 50% of swap-shuffled surrogate datasets for the other behavioral context. Based on these criteria, 12,408 3-neuron combinations were specifically enriched during social interaction, and 9,572 were specifically enriched during home cage exploration. There were 55,696 instances in which a social and nonsocial 3-neuron combination overlapped in 2 out of 3 neurons. In 97.0% of these cases, the neuron which was part of a social 3-neuron combination (triplet) but left out of the overlapping home cage triplet was part of a different 3-neuron combination that was enriched during homecage exploration (**Figure 6**, top right). Conversely, the neuron which was part of a nonsocial triplet but left out of the overlapping social 3-neuron combination was part of a different socially-enriched 3-neuron combination in 99.1% of cases (**Figure 6**, bottom right). Overall, an average of 71 enriched homecage combinations contained the neuron missing from the social triplet, and 85 enriched social combinations contained the neuron missing from homecage triplets. Thus, the specificity of a combination of co-active neurons for social vs. nonsocial behavior does not occur simply because some neurons were only active during one condition, but rather reflects the dynamic reorganization of patterns formed by neurons which are active in both conditions, i.e., changes in correlations. This – the behaviorally-specific occurrence of multineuron patterns of coactivity – represents the substrate through which correlations can add to the behavioral information transmitted by neuronal ensembles.

**Figure 6.**
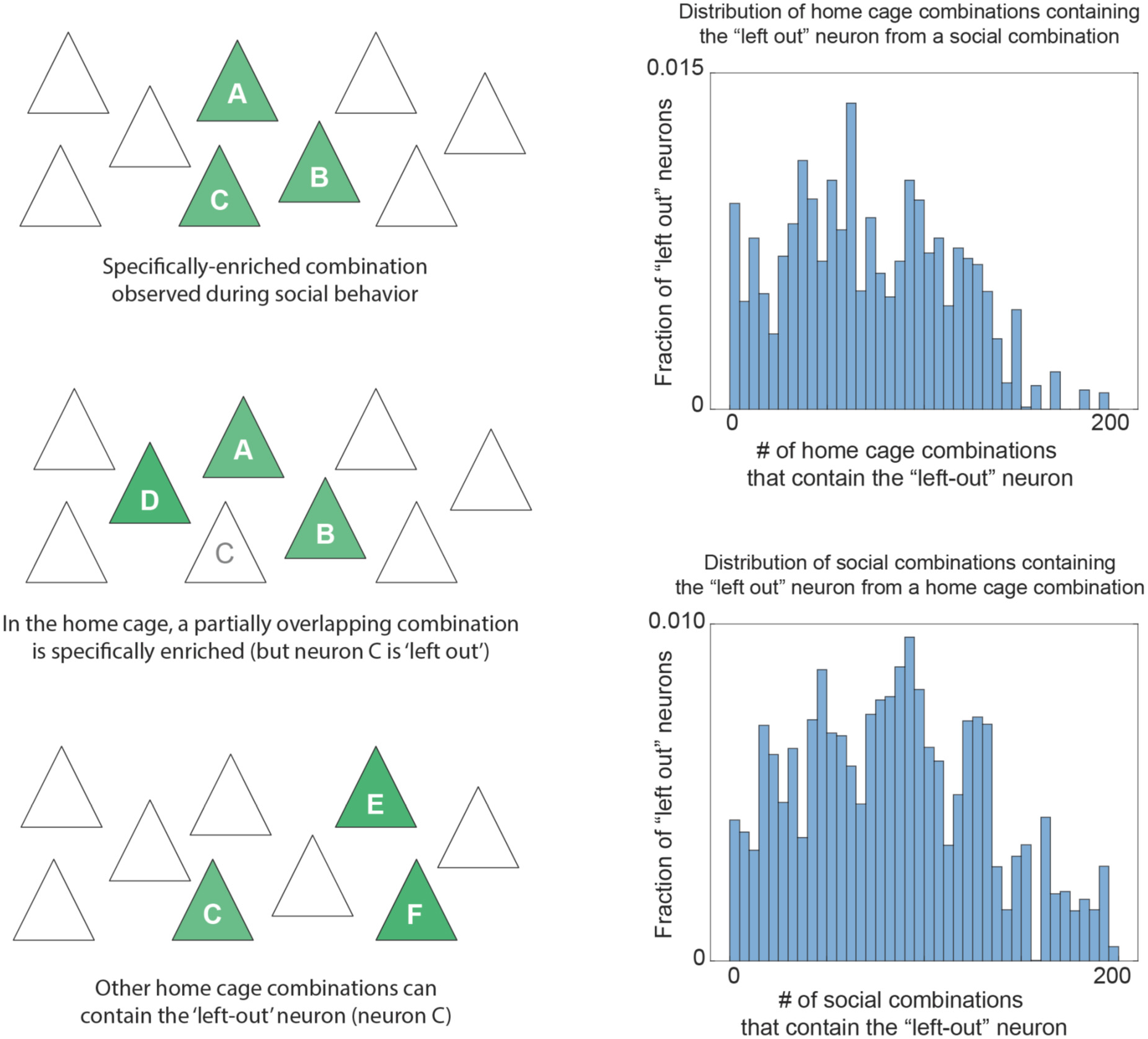
Behaviorally-specific patterns of coactivity are formed by neurons that are active in both conditions, but coactive with different partners in each condition. **Left:** We identified combinations of 3 neurons that are specifically enriched during one behavioral condition (occurring more often during social interaction than in 95% of surrogate datasets, and occurring less often during home cage exploration than in 50% of surrogate datasets, or vice-versa). We then identified overlapping combinations occurring during the opposite behavioral condition in which a single neuron was ‘left out.’ In other words, we identified combinations from the two conditions that overlapped in exactly two neurons. **Right, top:** Histogram showing the number of distinct 3 neuron home cage combinations that contain the neuron which participates in a social combination but is ‘left out’ during home cage behavior. **Right, bottom:** Histogram showing the number of distinct 3 neuron social combinations that contain the neuron which participates in a home cage combination but is ‘left out’ during social interaction. In the vast majority of cases, neurons that are ‘left out’ in one condition are still active during that condition and participate in other combinations.

### Socially-enriched combinations are deficient in Shank3 KO mice

We were curious whether there might be conditions under which these phenomena – the occurrence of multineuron combinations of coactivity during social behavior, and the ability of correlated activity to enhance the transmission of information about social behavior – might be impaired. To explore this, we performed microendoscopic GCaMP imaging in mice lacking the autism-associated gene Shank3 (23–25). These mice have been extensively studied as models of Phelan-McDermid syndrome, which often includes autism as a clinical feature. Shank3^−/−^ (KO) mice are known to have social deficits, and indeed, we found that compared to wild-type (WT) littermates, they spend significantly less time interacting with novel juvenile mice (**Figure 7A**).

**Figure 7.**
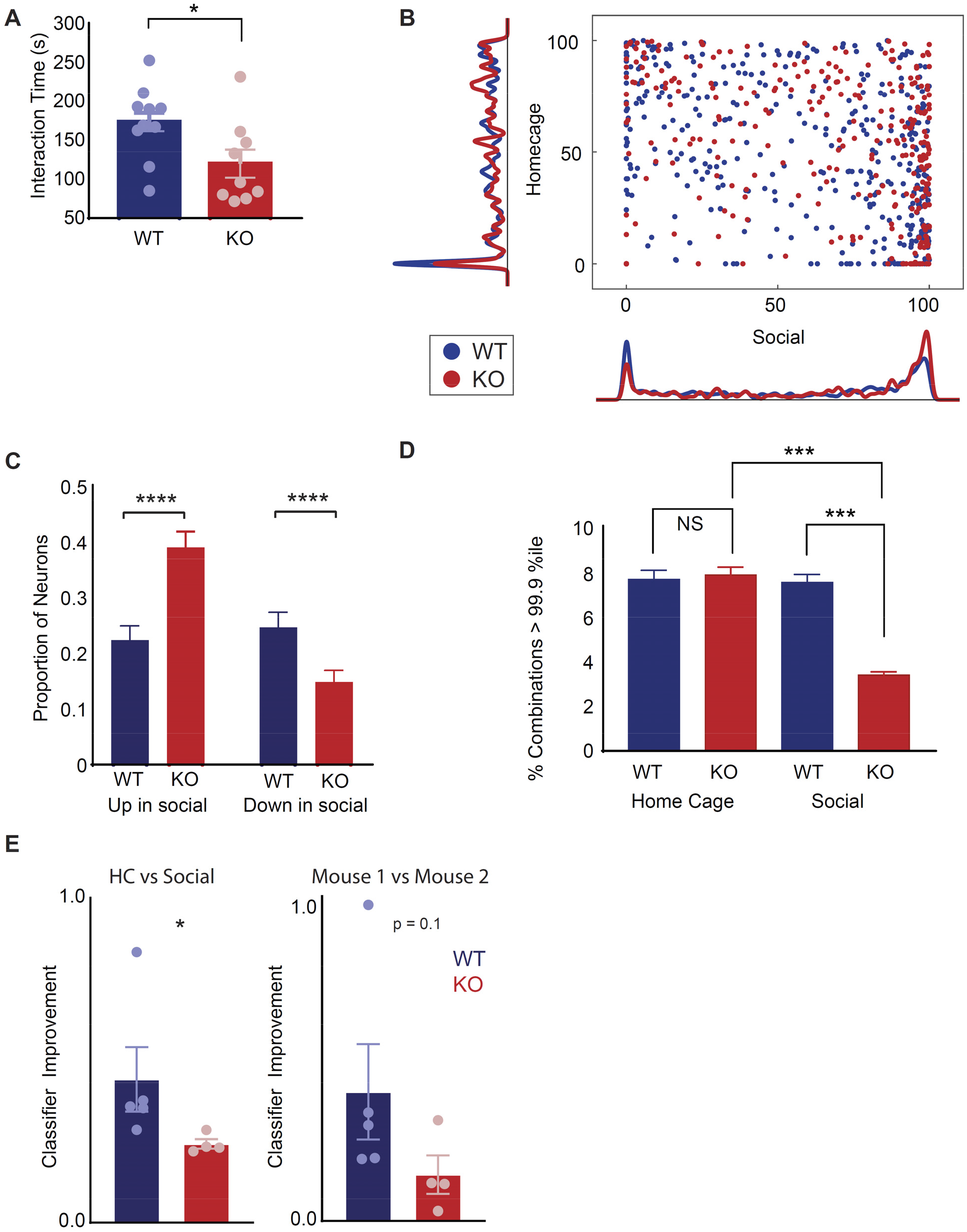
Shank3 KO mice have disorganized ensembles for which correlations fail to enhance the transmission of information about social behavior. **A.** The mean time that *Shank3* KO mice or wild-type (WT) littermates spend interacting with a novel juvenile mouse of the same sex during a 5 min assay. Data has been pooled from 8 unimplanted WT mice as well as the 5 implanted WT mice used for microendoscopic imaging, and 5 unimplanted KO mice in addition to the 4 implanted mice used for imaging. For implanted mice we used the average of interaction time for the 2 novel mice. Pooled data showed decreased interaction in KO mice (173 +/− 12 s vs. 120 +/− 18 s for WT and KO respectively, p < 0.05, t-test). The un-implanted cohort alone shows a similar significant decrease in interaction time for KO mice (165 +/− 15 s vs 110 +/− 16 s for WT and KO respectively, p < 0.05, t-test). In the implanted cohort there was a similar trend toward decreased interaction for KO mice (186 +/− 20 s vs 133 +/− 37 s, for WT and KO respectively, p = 0.21, t-test). **B.** (Similar to Fig. 1D). Scatter-plot showing the activity of each neuron during each behavioral condition, expressed as a percentile relative to a null distribution generated by circularly shuffling that neuron’s activity. Activity levels during social interaction or while the mouse was alone in its home cage are plotted on the x and y axis, respectively. Kernel density plots along the axes indicate the fraction of neurons whose activity was at a given percentile of the null distribution. Neurons with activity > 90^th^ percentile of shuffled datasets (green dotted line) were considered to be positively modulated, whereas neurons with activity < 10^th^ percentile (green dotted line) were considered to be negatively modulated during each behavior. Data is plotted for Shank3 KO mice (red) and WT littermates (blue) (mean percentile of activity during social interaction, WT: 50 +/− 2 percentile; KO: 64 +/− 2 percentile; p < 0.0001 by 2-sample *t*-test; mean percentile of activity during home cage, WT: 47 +/− 2 percentile; KO: 51 +/− 2 percentile; p = 0.1, *t*-test). **C.** Bar graph showing the fraction of neurons whose activity was positively or negatively modulated (>90^th^ percentile or <10^th^ percentile) during social interaction. The proportion of neurons which increased activity during social interaction was significantly greater in KO mice (22% in WT vs. 39% in KO, chi-squared = 17.7, p < 0.0001), whereas the downregulated ensemble was significantly smaller in KO mice (25% in WT vs. 15% in KO, chi-squared test, 8.2, p < 0.005). Error bars denote the binomial S.E.M. algebraically derived from total number of neurons and the proportion that were modulated in the specified direction. **D.** The proportion of 3 neuron combinations occurring during home cage exploration that are enriched > the 99.9^th^ percentile compared to swap-shuffled datasets was similar across WT (Blue; 7.6%) and KO (Red; 7.8%) mice. By contrast, the proportion of 3 neuron combinations occurring during social interaction that are enriched > 99.9^th^ percentile compared to swap-shuffled datasets was 7.5% in WT compared to only 3.4% in KO mice (total number of home cage combinations: 4,187 in 5 WT mice, and 5,878 in 4 KO mice; total number of social combinations: 5,487 in 5 WT mice, 16,326 in 4 KO mice). The top two plots show histograms of enrichment for the home cage (upper) or social conditions (middle); the lower panel is a bar graph showing the fraction of these combinations that were specifically enriched above the 99.9^th^ percentile (chi-squared = 165, p < 0.0001). Error bars denote the S.E.M. algebraically derived from the binomial distribution, the number of 3 neuron combinations in each condition, and the proportion of those combinations that were enriched. **E.** Bar graph showing how much better classifiers trained on real datasets perform whenb tested on SHARC shuffled datasets, compared to their performance when tested on swap-shuffled datasets, i.e. (***PerformanceSHARC - PerformanceSwap***)/(***PerformanceSwap - 0.5***). Each datapoint represents a classifier from one mouse. Left: The increase in performance was significantly larger in WT mice (blue) than KO mice (right) for classification of home cage periods (HC) vs. epochs of social interaction (Social), WT: 0.44 +/− 0.10 vs KO: 0.24 +/− 0.01, p = 0.016, Mann Whitney. Right: The increase in performance was non-significantly larger in WT mice (blue) than KO mice (right) for classification of interactions with Mouse 1 vs. those with Mouse 2, WT: 0.41 +/− 0.15 vs. KO: 0.14 +/− 0.06, p = 0.11, Mann Whitney. Combined analysis of both sets of classifiers (HC vs. Soc and Mouse 1 vs. 2) shows a significant effect of genotype: p < 0.05 by 2-way ANOVA.

We compared patterns of prefrontal activity in Shank3 KO mice and their WT littermates. As in WT mice, in Shank3 KO mice, many prefrontal neurons either increase or decrease activity during social interaction. However, compared to WT mice, the fraction of neurons whose activity increases during social interaction was significantly higher, whereas the fraction whose activity decreases was significantly lower (**Figure 7B-C;** 22% of 260 WT neurons vs. 39% of 290 KO neurons increased activity above the 90^th^ percentile of shuffled data during social interaction, chi squared = 17.7, p < 0.0001; 25% of WT vs. 15% of KO neurons decreased activity below the 10^th^ percentile of shuffled data during social interaction, chi squared = 8.2, p < 0.0001). Thus, Shank3 KO mice recruit abnormal neuronal ensembles during social behavior. We hypothesized that this might reflect a network-level disorganization that affects the normal patterning of activity during social behavior.

Indeed, we found that in KO mice a significantly smaller fraction of the 3-neuron combinations observed during social interaction were strongly enriched, i.e., occur more often in actual data than in 99.9% of swap-shuffled surrogate datasets (**Figure 7D**). This suggests that even though more neurons (i.e., larger ensembles), were recruited during social behavior in KO mice, these may have been less well-organized, such that the occurrence of socially-enriched patterns of activity is obscured by ‘noise,’ i.e., patterns formed by the chance overlap of activity between neurons that fire in a largely independent fashion. Notably, this deficiency was specific for social interaction. The fraction of 3-neuron combinations that were strongly enriched during home cage exploration (in comparison to swap-shuffled surrogate datasets) was similar in WT and KO mice (Figure 7D).

### The temporal organization of multineuron activity does not enhance the transmission of information about social behavior in Shank3 KO mice

The preceding shows that even though social behavior robustly recruits neuronal ensembles in Shank3 KO mice, the organization of these ensembles into multineuron combinations is disorganized. This suggests that the ability of patterns of correlated activity to encode information about social behavior may be impaired in these mice. To test this, we examined whether correlated activity contributes to the transmission of information about social behavior in Shank3 KO mice. As before, we generated swap and SHARC shuffled surrogate datasets, then tested the ability of classifiers trained on the original datasets (from Shank3 KO mice) to classify activity associated with behavior during social interaction vs. in home cage. We then compared the degree to which classifier accuracy was improved when testing SHARC-shuffled surrogate datasets compared to testing swap-shuffled ones. In datasets from Shank3 KO mice, we observed a significantly diminished improvement in classifier accuracy for SHARC vs. swap shuffled datasets, indicating a diminished role for correlated activity in encoding (relative improvement for classifying home cage vs. social: 0.44 +/− 0.10 in WT vs 0.24 +/− 0.01 in KO; relative improvement for classifying whether the subject was interacting with Mouse 1 vs. Mouse 2: 0.41 +/− 0.15 in WT vs. 0.14 +/− 0.06 in KO; 2-way ANOVA using classification type and genotype as factors: significant for genotype, p < 0.05, no effect of classification type, p = 0.58, or interaction, p = 0.76). Thus, in Shank3 KO mice, the multineuron patterns of coactivity which normally occur during social behavior are disturbed, and as a result patterns of correlated activity contribute less information about social behavior.

## DISCUSSION

During complex behaviors, the brain can use many strategies to represent information about the external environment and internal state of the organism. The term ‘ensemble’ is often used to refer to a group of neurons whose activity is similarly modulated (either increased or decreased) during specific behaviors (1,26–30). It is generally accepted that ensembles transmit behavioral information via changes in the activity levels of their constituent neurons. On the other hand, many studies have also shown that correlations between neurons can change during specific behaviors (3,10) or behavioral states (31–33). Importantly, correlations reflect changes in coactivity which exceed those expected to occur simply because of changes in the activity levels of the individual neurons (6). I.e., when an ensemble becomes more active, correlations between the neurons in that ensemble could go up, down, or remain unchanged. By optimizing synaptic interactions such as temporal summation, changes in correlated activity could potentially act to enhance the behavioral information transmitted by changes in ensemble activity, or transmit additional information.

The role of correlations in information transmission has been studied extensively in the isolated retina (34). In the cortex, noise correlations – correlations in trial-to-trial variability across neurons that are thought to be unrelated to the variable of interest – have also been extensively studied(11,13,14). Noise correlations can enhance information transmission, presumably by helping to disambiguate changes in activity levels due to the variable of interest from those attributable to other causes (noise). However, the ability of correlations to transmit information themselves, by changing as a function of the variable being encoded, has been less well studied. In particular, it is not clear whether changes in correlations encode additional information, beyond what is transmitted by changes in activity levels, and if so whether this reflects the behaviorally-specific occurrence of patterns of correlated activity / coactivity.

Here, we addressed this question using microendoscopic GCaMP imaging to measure activity from many (~40-100) prefrontal neurons during social behavior in mice. We used a simple neural network, in which prefrontal neurons provide input, there is one hidden layer, and a single output unit classifies social vs. nonsocial behavior, to quantify how well prefrontal ensembles would transmit information about social behavior to downstream neurons. We used multiple new approaches to disentangle the respective contributions of changes in activity levels vs. correlated activity to the encoding of social behavior. First, whereas previous studies have compared information transmission by real datasets vs. by shuffled ones(14), we extended a method we previously published, (22), to non-randomly shuffle datasets in order to preserve both behaviorally-modulated correlations and ensemble activity. This enabled us to compare the amount of information about social behavior transmitted by either SHARC-shuffled surrogate datasets or randomly-shuffled surrogates which preserved ensemble activity but not correlations. In this way, we found that correlated activity enhances the amount of information that prefrontal ensembles transmit about social behavior. Notably, we used neural networks to classify periods when a subject mouse was alone in its home cage vs. interacting with another mouse, and separate classifiers to determine whether the subject was interacting with test mouse 1 vs. test mouse 2. We found that correlated activity normally enhances the encoding of both types of information.

Second, when we examined connections within neural network classifiers, we found that prefrontal neurons which serve to detect social behavior increase their correlations during social behavior (whereas neurons which detect nonsocial behavior do not). Third, we found that combinations of coactive neurons which occur more often than expected based on the activity levels of the constituent neurons, manifest in a behaviorally-specific manner. Positive correlations measure neuronal coactivity that occurs more often than expected based on the chance overlap of activity between neurons. Thus, in accordance with our finding that social behavior modulates correlations, we found that multineuron patterns of coactivity which occur more often than expected by chance are behaviorally-specific. We also directly examined these behaviorally-specific and statistically-enriched combinations of coactive neurons. We found that they tend to be composed of neurons which are active in both conditions but only coactive in one, rather that neurons which are only active in one condition. Thus, we used multiple new approaches (comparing classifier accuracy for SHARC vs. swap shuffled datasets, measuring changes in correlations for neurons in ensembles derived from neural network classifiers, linking statistically enriched patterns of coactivity to specific behaviors) to show exactly how dynamic correlations encode information about social behavior.

Interestingly, statistically-enriched patterns of coactivity were specifically deficient during social behavior in mice lacking the autism-associated gene Shank3. Accordingly, in Shank3 KO mice, surrogate datasets which preserve behaviorally-modulated correlations failed to transmit more information about social behavior compared to randomly shuffled datasets which only preserved ensemble activity. This shows that the ability of correlated activity to enhance the transmission of information about social behavior is not automatic, and can in fact be disrupted in pathological states.

Similar to other recent studies (12,13), we have studied activity using binary activity rasters derived from GCaMP imaging. However, an important caveat is that any method of quantifying neural activity has limitations, such that there could be additional ways that neurons encode information which are not well resolved using this approach.

### What is the meaningful size of ensembles in the cortex?

Complex behavior is possible because the brain reliably encodes features pertaining to the external environment as well as the internal state of the organism. These features may be encoded by neuronal ensembles (1,26–30). What size of ensemble reliably encodes an aspect of behavior? We explored this question, by asking what input connection probability would optimize the ability of a downstream network to classify behavior based on input from prefrontal ensembles. Note: input connections in neural network classifiers do not necessarily correspond to actual connections in the brain – rather they provide information about the size and nature of neuronal ensembles across which information should be combined to most efficiently decode behavior. Peak classifier performance occurred for connection probabilities ~0.2 - 0.3. Performance was markedly lower when the connection probability was 0.5. This is surprising because a connection probability of 0.5 would maximize the entropy associated with each input connection. Thus, from the standpoint of encoding social behavior, combining activity from 20-30% of the input neurons may achieve some synergy that becomes degraded when ensembles are enlarged beyond this size. In the brain, nonrandom network connectivity (35,36) may similarly produce correlated activity / coactivity within ensembles of this size (37,38).

### Combinatorial codes vs. sequential patterns of activity

Like many recent studies, we measured population-level activity in the mouse neocortex using genetically encoded calcium indicators. These indicators transduce neuronal spiking on timescales ~100 msec. Thus correlated activity / ‘coactivity’ imply that neurons jointly increase their activity within windows ~100 msec, and do not necessarily imply synchronous spiking on faster timescales (milliseconds or even tens of miliseconds). At the same time, correlated activity / coactivity on these timescales should be differentiated from sequential activity of neurons observed during the performance of sequential behaviors (i.e. spatial navigation or overtrained tasks) in which the activity of specific neurons corresponds to moving through a specific location or performing a specific portion of a complex task. As discussed above, in the neocortex correlations and coactivity likely reflect recurrent neural network connectivity (37). By contrast, sequential patterns of neuronal activation can occur simply as a byproduct of the arrangement of spatial locations along a trajectory, the stereotyped order in which cues are encountered during a task, etc.

### Relevance to disease states

Interestingly, in *Shank3* KO mice, which exhibit social deficits, the mPFC successfully recruits specific neuronal ensembles during social interaction. However these ensembles are enlarged, their organization into statistically-enriched patterns of coactivity is disrupted, and correlations between neurons fail to enhance the information that these ensembles transmit. Thus, the computational units by which information is processed in the mPFC appears to be inefficient, i.e., social behavioral recruits an abnormally large number of neurons at the expense of the precise temporal patterning of this activity. This central finding is similar to other findings in rodent models of autism at both the single neuron and network levels (19,23,39). In particular, we found an increase in the recruitment of prefrontal neurons during social interaction. This mirrors a recent study which found hyperdynamic response to whisker stimulation in the same mice (23), possibly reflecting GABAergic circuit dysfunction and/or homeostatic compensations (40). (Note: these findings cannot be ascribed simply to the fact that Shank3 KO mice spend less time engaged in social interaction than their wild-type littermates; reduced interaction time would tend to reduce statistical power and thereby reduce the number of neurons that change their activity more than expected by chance.)

Increased excitatory activity causing decreased signal-to-noise ratio (SNR) has long been posited to contribute to the pathophysiology of autism (41). However the exact nature of ‘signal’ and ‘noise’, and the specific mechanism through which excessive activity degrades the SNR have been unclear. Here, we show how enlarged neural ensemble recruitment by specific behavioral conditions disrupts information transmission by degrading the ratio between statistically meaningful patterns of coactivity (the signal) and the random overlap of activity between neurons (noise).

## MATERIALS AND METHODS

### Behavior

C57/B6 mice were obtained from Jackson Laboratories. We utilized adult mice of either sex housed and bred in the UCSF animal facility. Adult mice were habituated to the room and observer for 3 days prior to test day. All videos were subsequently scored by a blinded observer. For imaging experiments, 5 WT and 4 KO littermates were generated through crosses between Shank3 heterozygous parents and injected with AAV5.Syn.GCaMP6f.WPRE.SV40. We included an additional 5 WT mice which were injected with AAV5.Syn.GCaMP6m.WPRE.SV40 (42). Viruses were obtained from Penn Viral Core. Injections and 500 um GRIN lens (Inscopix) implantations were carried out in 8-12 week old mice to express GCamp6f in prefrontal cortical neurons under control of the human Synapsin promotor. Mice were anesthetized with 2% isoflurane and mounted in a stereotactic frame. Craniotomies were made according to stereotaxic coordinates relative to Bregma. Coordinates for injection into mPFC were (in mm relative to Bregma): +1.7 anterior–posterior (AP), –0.3 mediolateral (ML) and –2.75 dorsoventral (DV), and GRIN lenses were implanted at the same AP and ML coordinates, to a depth of 2.25. We subsequently attached baseplates for attaching the microendoscope, ~4 weeks later depending on GCamp expression. Mice were habituated for three days with the scope attached, prior to test day. On test day, mice were habituated with the scope turned on, then imaged in alternating home cage and social epochs. During social epochs, one of 2 novel sex-matched juvenile mouse was introduced to the test mouse’s homecage, in sequential order so that there were two ‘novel’ epochs, followed by two ‘familiar’ epochs interleaved with ‘home cage’ epochs during which the juvenile mice were removed and the test mouse was free to explore its home cage. The first and last home cage epoch were 10 minutes in length; the others were 5 minutes in length. Each social epoch lasted 10 minutes but only the first 5 minutes were recorded and scored. During each behavioral epoch, observer was not in the room. Interaction epochs were defined from the moment test mouse first sniffed the juvenile conspecific or object, until the test mouse turned away. Videos were recorded using Anymaze, and scored by a blinded observer. For the bulk of analysis we pooled data across 10 WT mice. Shank3 KO mice were compared only to recordings from WT littermates.

### Image acquisition and segmentation

Images were acquired using an Inscopix nVoke micreoendoscope attached to a laptop computer and synced to a separate video acquisition computer running Anymaze. Frame rate was 20 Hz and the laser power was 0.2 mW. Acquisition was performed using 2×2 pixel binning, then subsequently downsampled again by 2.

We segmented neuronal signals using a modified PCA/ICA approach (20,21), modified so that each segment was expressed as a binary ROI consisting of pixels representing a single neuron. I.e., we used the output from the PCA/ICA to identify a set of contiguous pixels which represent a neuron, then averaged fluorescence signals across those pixels. To deconvolve neuronal signals from background neuropil signals, we also subtracted the mean signal from each identified segment from the mean value in pixels surrounding the edge of the segment (we excluded pixels that belonged to another ROI). Signals were subsequently lowpass filtered to remove high frequency noise using the Matlab command: designfilt(“lowpassfir”, “PassbandFrequency”, 0.5, “StopbandFrequency”, .65, “PassbandRipple”, 1, “StopbandAttenuation”, 25). All signal traces shown represent normalized versions of the *dF*/*F*_*0*_ trace, where *F*_*0*_ *is estimated by the median* value in the surround region. Threshold based event detection was performed on the traces by detecting increases in *dF*/*F*_*0*_ exceeding 3σ over one second, then only keeping those events which exceeded a 15σ increase over two seconds, and a total area under the curve of 250σ. As there were occasional downward deflections due to surround subtraction, we instituted a final parameter requiring that the peak cross an absolute value of *dF*/*F*_*0*_ = 0.0125. σ is the standard deviation of *dF*/*F*_*0*_, calculated over the least active 50% of the movie. In some cases these parameters were adjusted slightly to optimize event detection to > 95% sensitivity and specificity, based on visual inspection, for each movie. After identifying these events in the GCaMP signal from a cell, the cell was considered “active” during the entire period from the beginning of an event until the GCaMP signal decreased 30% from the peak of the event, up to a maximum of 2 seconds. The peak of the event was defined as the local maximum of the entire event, from the beginning of the event until *dF*/*F*_*0*_ returned to the pre-event baseline value. Calcium traces from segmented neurons were visually inspected and a small number of segments were removed if they did not appear to represent a single, unduplicated neuron. We restricted further analysis to those mice with 25 or more active neurons. We then created 2-dimensional event rasters consisting of detected events for each neuron over the course of the experiment.

### Detection of behaviorally modulated neurons

To determine the response of individual neurons to behavioral context, we averaged the activity of each neuron during frames corresponding to periods of social interaction, or to a temporally matched set of frames during the preceding home cage epoch. We then created a ‘null distribution’ for each neuron that represents the percent of time active expected in each condition based on chance, by circularly shuffling the data 10,000 times. We then compared the activity of each neuron during either social interaction or home cage exploration to this null distribution. Neurons were considered positively modulated if they exceeded the 90^th^ percentile of that observed in circularly-shuffled datasets, and negatively modulated if the percent of frames that a neuron was active during a given context was below the 10^th^ percentile of observations from circularly-shuffled data.

### SHARC

SHARC (<u>SH</u>uffling <u>A</u>ctivity to <u>R</u>earrange <u>C</u>orrelations) is an iterative method for generating surrogate datasets. SHARC nonrandomly shuffles blocks of activity within a raster to generate a new (surrogate) raster in which the pairwise correlations between neurons match a target correlation matrix (21). Here we apply this previously-published method, with modifications to also preserve the activity level in each neuron (Figure 4B).

To begin, note that each raster is equivalent to a collection of blocks of activity. Each block of activity is defined by the time at which it begins, its duration, and the neuron which is active. On each iteration one block of activity is randomly chosen and assigned to a new neuron as follows. Suppose block *i* has been chosen to be re-assigned. First, we find all the blocks of activity that overlap with block *i*. Next, we selected the subset of these blocks for which new cell identities had already been assigned. Call this set *X*. Let *r*_*j*_ represent the number of timepoints over which block *j* ∈ *X* overlaps with block *i*, and let *n*_*j*_ represent the identity of the cell assigned to block *j* ∈ *X*. *L*_*i*_ and *L*_*j*_ are the lengths of blocks *i* and *j*, respectively. Then we constructed a vector,

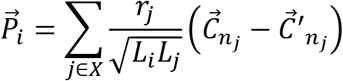

where 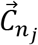 represents row *j* of the target correlation matrix, i.e. the target correlations between neuron *n*_*j*_ and the other neurons, and 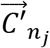 contains the current values of the correlations between neuron *n*_*j*_ and the other neurons based on the blocks of activity that have already been re-assigned. This step can be thought of as “guessing” which cell *should* be assigned to a particular block of activity by first figuring out what other cells are active at the same time, then choosing cells which are strongly correlated with these known active cells. Note that we assign values of 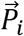 (i.e., construct “guesses” about which cell should be active), using the *difference* between the current correlation matrix 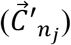 and the target correlation matrix 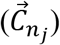, in order to identify cell pairs for which the current correlation deviates from the target value, and force the new correlation matrix to progressively approximately the target correlation matrix.

We set elements of 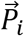 to zero if the corresponding neuron had already been assigned to a block of activity that overlaps with block *i*, i.e. element *n*_*j*_ of 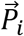 was set to zero ∀ *j* ∈ *X*. Finally, we assigned block *i* to the neuron corresponding to the maximum value of 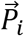. This can be thought of as choosing the cell that represents the “consensus” based on tallying up all of the “guesses” about which cells “should” be assigned to the block of activity being considered.

When all the elements of 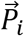 were zero, e.g. because there no overlapping blocks of activity have had new cell identities assigned yet, then we chose a cell in order to match the originally observed level of activity. Specifically, after every iteration, we kept a log of the net number of blocks of activity that each neuron had donated or received. We used this vector to create a weighted probability whereby events from neurons which had received a net positive number of blocks were more likely to be chosen to be reassigned. To further ensure that the total number of active events for each neuron in the surrogate dataset was similar to the real dataset, if the difference between the number of blocks gained – lost in the reassignment process exceeded +4 for a particular neuron, then that neurons was no longer eligible to receive additional blocks of activity; similarly blocks of activity belonging to neurons who had lost greater than 3 blocks of activity were no longer eligible be reassigned. As the selection of blocks of activity for reassignment is probabilistic, we first perform a swap shuffle to ensure that all blocks of activity in the original dataset are swapped at least one time, then perform a total of 5 reassignments for each block of activity in the source dataset.

We extended this approach to generate surrogate datasets by shuffling data within shorter time windows (i.e., individual behavioral epochs). Here a discrete set of frames is chosen, corresponding to a subraster of the original raster. By repeating the process described above for each subrasters, then recombining the shuffled subrasters, we generate a complete shuffled dataset.

### Classifier

We designed and trained a neural network to classify behavior (periods when a mouse was alone in its home cage vs. engaged in social interaction). This network contained 1000 units in a hidden layer, each of which received input from specific prefrontal neurons (from the real dataset). Thus, in each frame the activity of each hidden layer unit was just the summed activity of the connected prefrontal neurons. Each hidden layer unit had an output weight that represented the strength of its connection to a single output unit. On each frame the activity of the output unit was computed as:

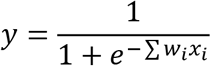

where *w*_*i*_ is the output weight from hidden unit *i* and *x*_*i*_ is the activity of hidden unit *i*.

When we performed training and testing using the same dataset, we divided the dataset into alternating blocks of 500 frames for training vs. testing (in other cases we used the real dataset for training, then tested using a surrogate dataset). We restricted training or testing to frames in which mice were scored as actively engaged in social interaction (or matched frames during periods when the mouse was alone in its home cage). We also limited training / testing to frames with at least 3 active neurons.

We trained the output weights by performing 500 passes through the training data (each pass visited all of the training frames in a random order). On each training timestep, we calculated *y*, the activity of the output unit, and then adjusted each output weight based on:

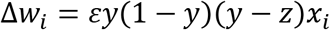

where z is the correct classification of the frame (0 for social behavior, 1 for home cage) and *ε*, the learning rate, was set to 0.05.

Following training, we examined the pattern of input connections and output weights. The distribution of output weights was roughly gaussian and centered near 0. We identified the selection of prefrontal neurons most likely to be connected to hidden layer units with large positive or negative weights. Hidden layer units with large negative or positive output weights bias classification towards the social or home cage condition, respectively. Therefore, we refer to the 25 hidden units with the most negative or positive weights as ‘social’ or ‘home cage’ units respectively. We calculated the number of input connections between each prefrontal neuron and the 25 home cage units or 25 social units. We then defined ‘home cage’ or ‘social’ ensembles as the 20% of prefrontal neurons with the most input connections to home cage or social units, respectively. As described in the main text, we then analyzed properties of these two ensembles.

### Quantification of multineuron combinations

Estimating chance overlap between activity of largely independent neurons requires accounting for two factors. First, neurons with higher activity are more likely to overlap by chance with other neurons. Second, overall network activity is dynamic over time, creating a tendency for otherwise independent neurons to be recruited at similar times. Thus, it is necessary to identify combinations which occur more often than expected based on 1) the activity levels of the constituent neurons, and 2) the fact that activity in a network is not constant over time. We can do this by quantifying the occurrence of combinations in datasets which have been shuffled to preserve 1) the overall level of activity in each neuron, and 2) the total level of activity in the network at each point in time.

3 neuron combinations were quantified by identifying each combination present in frames in which 2 or more neurons were active. The number of frames each combination was active in *real data* was stored in a n-dimensional matrix. Surrogate datasets were then generated from event rasters by *swapping* the identity of neurons associated with detected events (periods of activity). As the timing of events themselves is unchanged, and only the identity of the participating neurons are exchanged, this preserves both the number of events per frame and the number of events that each neuron participates in. Therefore, the total number of combinations in each frame and over the course of the experiment (i.e., the sum of occurrences across all combinations) is also preserved. The total number of combination occurrences in which a given neuron participates would also tend to be preserved in these swap-shuffled surrogate datasets.

We then quantified how often each combination occurred in real vs. swap-shuffled data. By comparing how often each combination occurred in real data vs. in 1,000 swap-shuffled surrogate dataset, we were able to quantify how ‘enriched’ each combination was, compared to the level of occurrence expected by chance based on the activity levels of its constituent neurons (and the overall temporal pattern of network activity). We expressed enrichment as a percentile, calculated relative to swap-shuffled surrogate data, e.g., the 100^th^ percentile indicates that a particular combination occurred more often in real data than in all 1,000 surrogate datasets. Further analysis was restricted to ‘enriched combinations’, i.e., combinations that occurred more often in real datasets than in 95% of surrogate datasets.

### Generation of Synthetic datasets

all synthetic datasets consisted of 100 neurons and 6000 frames with an overall activity A of 5%. Briefly we randomly chose n active neurons in each frame f where *n* = *A* + *sin*(*f*)*A*. Nonoverlapping patterns or assemblies each consisting of 8 neurons were inserted into this oscillating network reciprocally swapping activity from non-assembly neurons to assembly neurons in each frame in which the first neuron of the assembly/pattern was active. In frames in which less than 8 neurons were active we randomly chose neurons from the assembly to be activated. To maintain equivalent levels of activity reciprocal swaps were made by choosing another frame in which the assembly neuron receiving activity was active and reassigning activity in that second frame from the assembly neuron to the non-assembly neuron. To create datasets in which the activity of individual neurons was modulated between ‘State A’ and ‘State B’ we transferred a proportion of active events from half the neurons to partner neurons so that half the neurons increased their activity and half the neurons activity decreased. For each pair of neurons the proportion of transferred activity was a random number between 0 and the specified upper bounds of modulation which ranged between 5 and 50%. We trained our classifier on blocks consisting of 50% of each dataset and testing on the remainder.

### Statistical analysis

Neurons and significant combinations from all animals and groups were pooled and counted as single units. Proportions were compared using chi-squared test. Activity levels were compared using paired *t*-tests (2-sided), unless otherwise noted. Where applicable, error bars denote standard error. Values of the classifier performance (accuracy) were generated by averaging after re-running the training / testing procedure at least 25 times.

## Acknowledgments

We thank Josh Berke and Ruchi Malik for helpful comments on the manuscript.

## Data and materials availability

All code used in the analysis will be deposited on GITHUB for any researcher for purposes of reproducing or extending the analysis.

---

**Figure S1.**
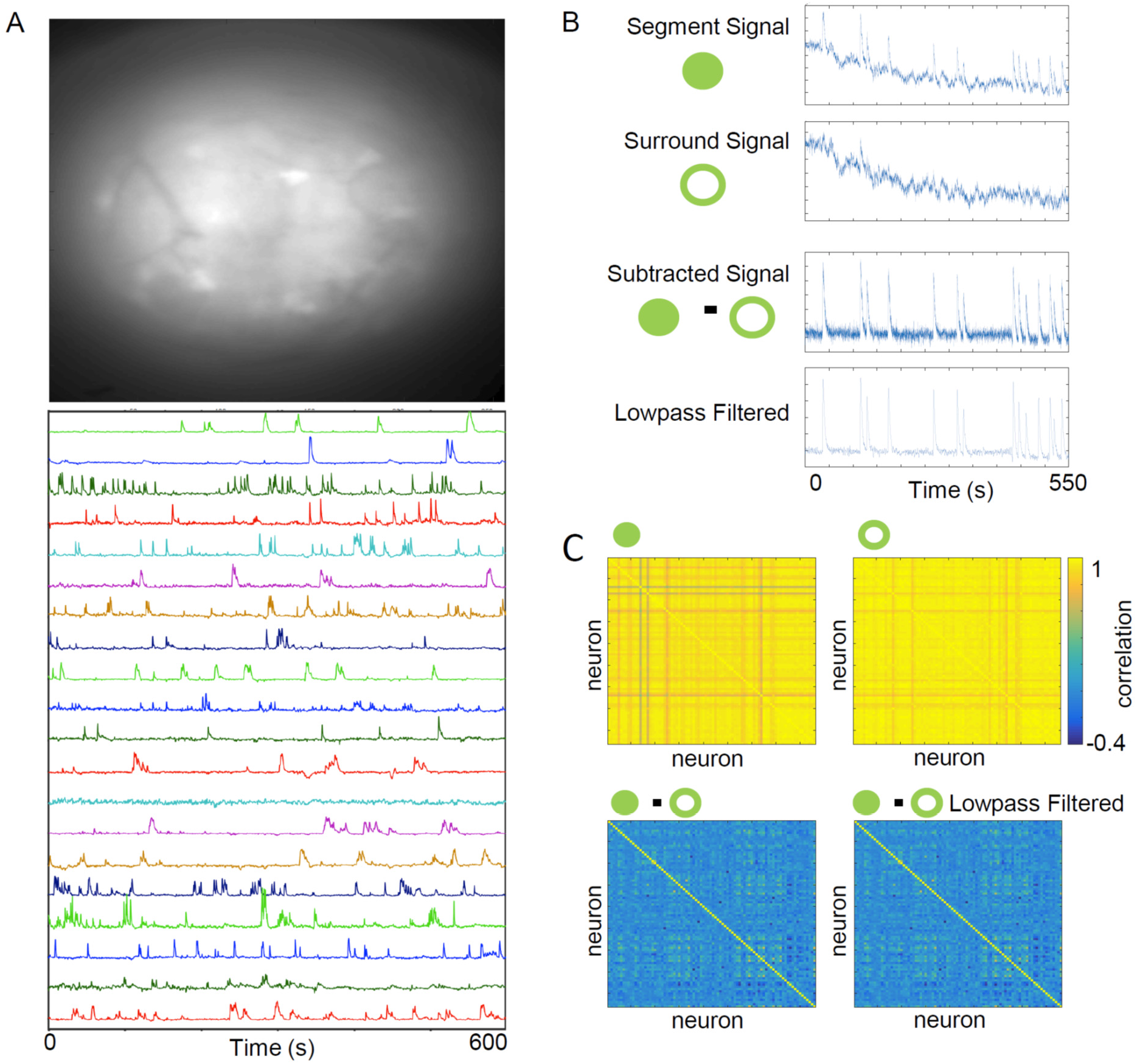
Spatial Decorrelation of Neuronal Signals. **A.** Example image (top) and individual neuron GCaMP traces (bottom) from prefrontal cortex imaged with implanted endoscope. **B.** The average GCaMP signal from a region of interest (ROI), corresponding to one neuron, was corrected by subtracting the average GCaMP signal from the surrounding pixels, in order to spatially deconvolve signals from each ROI vs. the surrounding neuropil. Examples traces from a single neuron are shown. **C.** The pairwise correlation matrix between signals from different neurons is shown (calculated from 550 seconds of activity from a single wildtype mouse), for the original GCaMP signals (top left), the surround signals (top right), the surround-subtracted signals (bottom left), and the surround-subtracted signals after lowpass filtering (bottom right).

**Figure S2.**
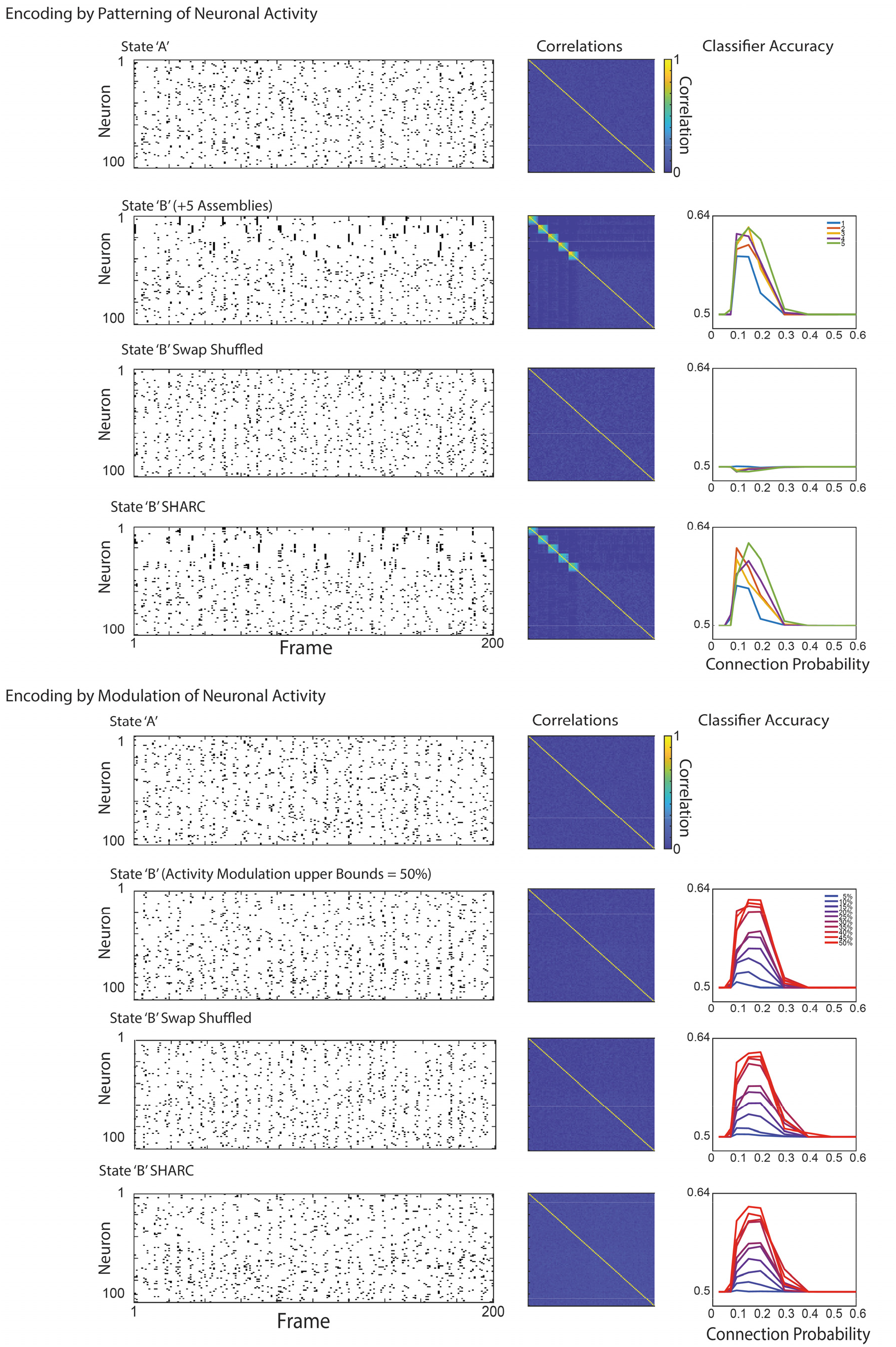
Patterns of correlations and the accuracy of classifications that are based on correlated activity are both intact after SHARC shuffling. **Top:** As described in the main text we created synthetic datasets comprising activity of 100 neurons during two states. The activity raster for ‘State B’ was generated by modifying the ‘State A’ raster. Specifically, in State B we ‘inserted’ between 1 and 5 cell assemblies (groups of neurons with correlated activity) by reciprocally swapping activity epochs so that both the overall level activity in each neuron, and total number of neurons active in each frame, were unchanged between the ‘State A’ and ‘State B’ rasters. <u>Left column</u>: the first 200 frames of activity for ‘State A’ and ‘State B’, followed by swap or SHARC-shuffled versions of the original ‘State B’ raster. <u>Middle column</u>: correlation matrices for raster. <u>Right column</u>: accuracy of a neural network trained to classify the original ‘State A’ and ‘State B’ rasters, tested using either the original ‘State B’ raster or swap or SHARC shuffled versions. Note that SHARC, but not swap shuffling, maintains both the pattern of correlations as well as the ability of the classifier to accurately decode ‘State A’ vs. ‘State B’. **Bottom:** Similar to Top, but now we created the ‘State B’ raster from the ‘State A’ raster by randomly shifting a fraction of activity from neurons 1-50 to neurons 51-100. Thus, ‘State A’ and ‘State B’ are differentiated by the activity levels of neurons. Note that in this case the classifier functions independently of correlations and is not affected by swap shuffling the ‘State B’ raster.

**Table S1.** Details for each mouse included in this study. The table shows the genotype, sex, number of imaged neurons, and peak classifier accuracy (performance when half the data was used for training and half for testing). Figure 2 showed how classifier accuracy depends on the input connection probability; because their performance was not >50% for multiple input connection probabilities, WT mice 3 & 5 (marked with #) were not included in this illustrative plot. However, we did not exclude data from these mice in any analyses. WT mice 1-5 were wild-type littermates of Shank3 KO mice. WT mice 6-10 (indicated by italics) were not littermates of Shank3 KO mice and were therefore not included as WT controls for the analyses shown in Figure 6.

| Mouse | Sex | Neurons | Peak Accuracy |
| --- | --- | --- | --- |
| WT |  |  |  |
| 1 | f | 84 | 0.687 |
| 2 | f | 39 | 0.705 |
| #3 | m | 25 | 0.53 |
| 4 | f | 75 | 0.605 |
| #5 | m | 37 | 0.459 |
| 6 | *m* | 86 | 0.647 |
| 7 | *m* | 74 | 0.848 |
| 8 | *m* | 73 | 0.691 |
| 9 | *m* | 71 | 0.694 |
| 10 | *f* | 99 | 0.633 |
| KO |  |  |  |
| 1 | f | 83 | 0.772 |
| 2 | m | 77 | 0.832 |
| 3 | f | 45 | 0.34 |
| 4 | f | 85 | 0.627 |

